# Analysis of the expression of PIWI-interacting RNAs during cardiac differentiation of human pluripotent stem cells

**DOI:** 10.1101/639906

**Authors:** Alejandro La Greca, María Agustina Scarafía, María Clara Hernández Cañás, Nelba Pérez, Sheila Castañeda, Carolina Colli, Alan Miqueas Möbbs, Natalia Lucía Santín Velazque, Gabriel Neiman, Ximena Garate, Cyntia Aban, Ariel Waisman, Lucía Moro, Gustavo Sevlever, Carlos Luzzani, Santiago Miriuka

## Abstract

PIWI-interacting RNAs (piRNAs) are a class of non-coding RNAs initially thought to be restricted almost exclusively to germ line cells. In recent years, accumulating evidence has demonstrated that piRNAs are actually expressed in somatic cells like pluripotent, neural, cardiac and even cancer cells. However, controversy still remains around the existence and function of somatic piRNAs. Using small RNA-seq samples from H9 pluripotent stem cells differentiated to mesoderm progenitors and cardiomyocytes we identified the expression of 447 piRNAs, of which 241 were detected in pluripotency, 218 in mesoderm and 171 in cardiac cells. The majority of them originated from the sense strand of protein coding and lncRNAs genes in all stages of differentiation, though no evidences for secondary piRNAs (ping-pong loop) were found. Genes hosting piRNAs in cardiac samples were related to critical biological processes in the heart, like contraction and cardiac muscle development. Our results indicate that somatic piRNAs might have a role in fine-tuning the expression of genes involved in the differentiation of pluripotent cells to cardiomyocytes.

## Introduction

Differentiation of pluripotent stem cells (PSC) to cardiomyocytes (CM) was first reported shortly after the characterization of embryonic stem cells (ESC) (*1*). Initially, differentiation was non-specific and spontaneously achieved, but in the last 10 years upgraded protocols have been developed significantly improving efficiency and reproducibility in cardiac differentiation (*2–4*). These protocols are based in sequentially adding factors (morphogens) and/or inhibitors that modulate Wnt/*β*-catenin signaling pathways in pluripotent cells. PSC-based models undergo epithelial-to-mesenchymal transition to an early mesoderm progenitor cell (MPC) (*2, 5*) followed by further committing to cardiac mesoderm and later cardiac progenitor cells (CPC), which may eventually adopt more especialized features. Though this is arguably similar to *in vivo* embryo development, they recapitulate hallmark features of differentiation thus becoming well suited tools for disease modelling, drug screening and potential cell-based therapies.

Like many other developmental processes, changes associated with differentiation to CM are tightly regulated. Only recently, and mostly due to the advent of next generation sequencing technologies, the scientific community is unveiling the complex regulatory networks governing the shifts in gene expression profiles. Non-coding RNAs (ncRNAs) are critical players in these networks, regulating almost all cellular processes including proliferation, differentiation and death (*6, 7*). Although microRNAs (miRNAs) are the most extensively studied in a wide variety of organisms (*8–11*), other ncRNAs have been identified such as long non-coding RNAs (lncRNAs), small interfering RNAs (siRNAs), circular RNAs (circRNAs) and PIWI-interacting RNAs (piRNAs). Much has been published about these ncRNAs, though piRNAs is one of the least understood. Thought to be initially confined almost exclusively to germinal cell lines (*12*), piRNAs gained much attention primarily because of an increasing amount of evidence demonstrating that these ncRNAs are not only expressed in somatic cells but they actively participate in gene regulation as well (*13–16*).

Expression of piRNAs was first described as negatively regulating transposition of repetitive elements thus protecting genome integrity and favouring selfrenewal (*12, 17*). Reports in numerous organisms showed that they exert their regulatory function through binding a specific clade of the Argonaute (AGO) family-namely PIWI proteins-, resulting in an association which resembles the well-known AGO/miRNA complex (*8, 18–20*). Unlike miRNAs, piRNAs are primarily biosynthesized as single-stranded long precursors which are then clived into the 24-34 nucleotide-long mature forms in a Dicer-independent manner (*12, 21, 22*). They show a bias for uridine (U) redidues in 5’ ends together with a 2’O-methyl modification at their 3’ ends. Germ line piRNAs were also found to be synthesized through a secondary pathway named the Ping-Pong amplification loop, which increases levels of primary piRNAs using target mRNAs as intermediary molecules for processing new piRNA precursors (*20, 23*). Of note, these mechanisms seem to be highly conserved across species (*18, 24*).

In the last few years, many studies have proposed an active participation of PIWI/piRNAs complexes in diverse and critical pathways such as neural development or body regeneration of lower eukaryotes (*25, 26*). Furthermore, recent work demonstrated a positive correlation between altered piRNA expression profiles and clinically relevant pathologies. The involvement of specific piRNAs in regulating mRNAs levels of genes related to Alzheimer’s disease was described in 2017 (*27*), while other groups implicated piRNAs in cardiac function and regeneration through modulation of AKT pathwway (*28, 29*). However, great controversy still remains around expression, function and biosynthetic pathways of somatic piRNAs. Particularly, the potential role of piRNAs in differentiation of pluripotent stem cells to cardiomyocytes has not been formally addressed. Using small RNAseq data generated in our laboratory (*11*) we characterized the expression profile of small RNAs consistent with piRNAs in three stages of cell differentiation from pluripotency (day 0) to mesoderm (day 3.5) and then contractile cardiocytes (day 21). Results presented here provide evidences supporting the existence of somatic piRNA transcripts and their stage-specific pattern as a mechanism for potentially fine-tuning gene expression during cell differentiation.

## Results

### Detection and characterization of piRNA

Detection of piRNAs was conducted on small RNAseq samples from three independent experiments consisting of pluripotent stem cells (PSC, day 0), early mesoderm progenitor cells (MPC, day 3.5) and cardiac progenitor cells (CPC, day 21). After aligning reads to human reference genome (hg38), we found that more than half of mapped reads were 20 to 23 nucleotides (nt) long, where the abundant miRNAs are included (Figure 1a, (*11*)). Considering that the average length of piRNAs in mammals ranges between 24 and 34 nt (*12, 22, 30*), mapped reads were filtered by length to accommodate to this restriction. Nearly 50-70% of mapped reads were removed from the samples after this initial processing step (Supplemental Figure 1a and Supplemental Table 1). Then, employing a similar approach as previously published work (*15*), we filtered out any read that mapped on ncRNAs besides piRNAs given that previous publications emphasized on the fact that many identified piRNAs were actually fragments of other types of ncRNAs (*31*). Approximately 5-20% of initial mapped reads remained after this step (Supplemental Table 1). Importantly, all nine aligned samples behaved similarly to both filtering steps (Figure 1b), reflecting consistency among experimental replicates.

**Figure 1:**
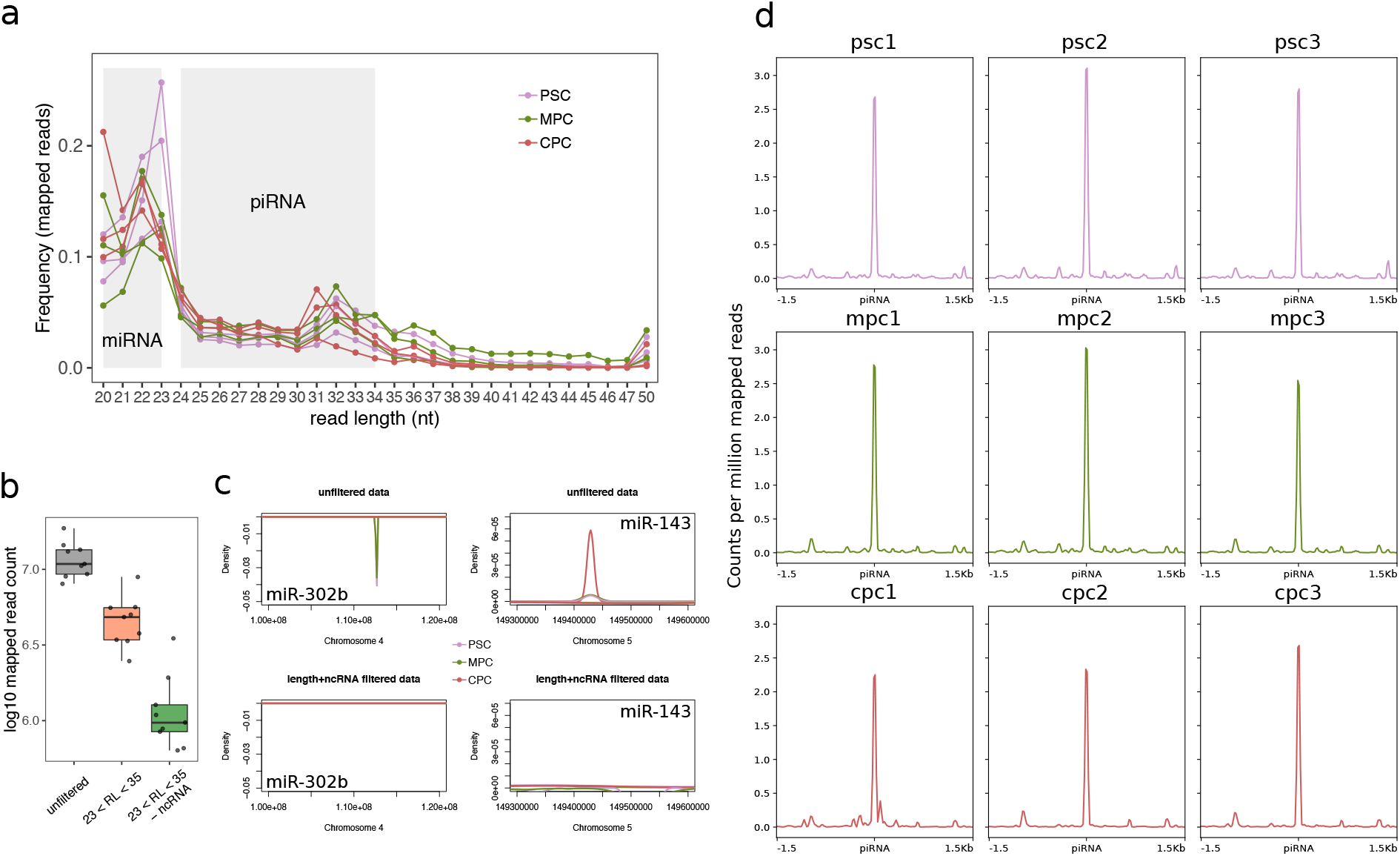
Detection of piRNAs in small RNAseq samples. a) Length frequency of unfiltered mapped reads in three samples from pluripotent cells (PSC), three from mesoderm progenitor cells (MPC) and three from cardiac cells (CPC). Grey areas denote the size range for both microRNAs (miRNA) and putative piwi-interacting RNAs (piRNA). The color key used is indicated in top right corner of the plot. b) Number of mapped reads for all nine samples before processing (unfiltered, grey box), after filtering by read length (23 < RL < 35, coral box) and removing non coding RNA other than piRNAs (−ncRNA, green box). c) Distribution of mapped reads as a function of density on a fragment of chromosome 4 (chr4:100,000,000-120,000,000; −) which includes pluripotency miR-302b and a fragment of chromosome 5 (chr5:149,300,000-149,600,000, +) including cardiac miR-143 for unprocessed alignments (top panels, unfiltered data) and fully processed samples (bottom panels, length+ncRNA filtered). Alignment files from each experimental replicate were merged into one. Color key for the density curves is shown in the graph. d) Analysis of coverage on all piRNA loci available in piRbase for fully processed normalized (counts per million) samples.

To verify the elimination of potential misleading contaminants in fully processed alignments, we analyzed the distribution of mapped reads over two well-characterized miRNAs, pluripotency-associated miR-302b and cardiac-expressed miR-143 (*11*). As expected, expression of miR-302b was evident in unfiltered data of PSC and MPC populations while miR-143 showed appreciable coverage in unfiltered data of CPC (Figure 1c, top panels). No signal was detected for either of the two genes in processed alignments (Figure 1c, bottom panels). However, these samples showed a strong and sharp coverage signal on known piRNA loci (Figure 1d) confirming that the pipeline employed successfully enriches for reads mapping to these known piRNAs. Henceforth, all analyses were performed on fully processed alignments unless explicitly specified otherwise.

Sequence analysis of reads mapping to known piRNA loci showed that all samples but MPC bore a bias for 5’ uridine residues as it usually occurs in germline cells (Figure 2a). We corroborated our proceedings by employing the pipeline described above on two small RNA-seq samples from human test is downloaded from the ENCODE project. Indeed, there was a marked preference for uridine at 5’ ends in testis samples (Supplemental Figure 2a and b), which suggest that putative piRNAs in our model are subjected to similar mechanisms of 5’ end formation as in germline cells. However, the substantial difference in frequecy of 5’-U residues between our samples and testis samples could be an indicative of unconserved biosynthetic steps (*26*). In addition, no secondary piRNA production was detected in any of the replicates of our samples given that we did not found evidences of the characteristic 10 nt overlap (ping-pong signature) between 5’ ends of sense and antisense mapped reads (Figure 2b).

**Figure 2:**
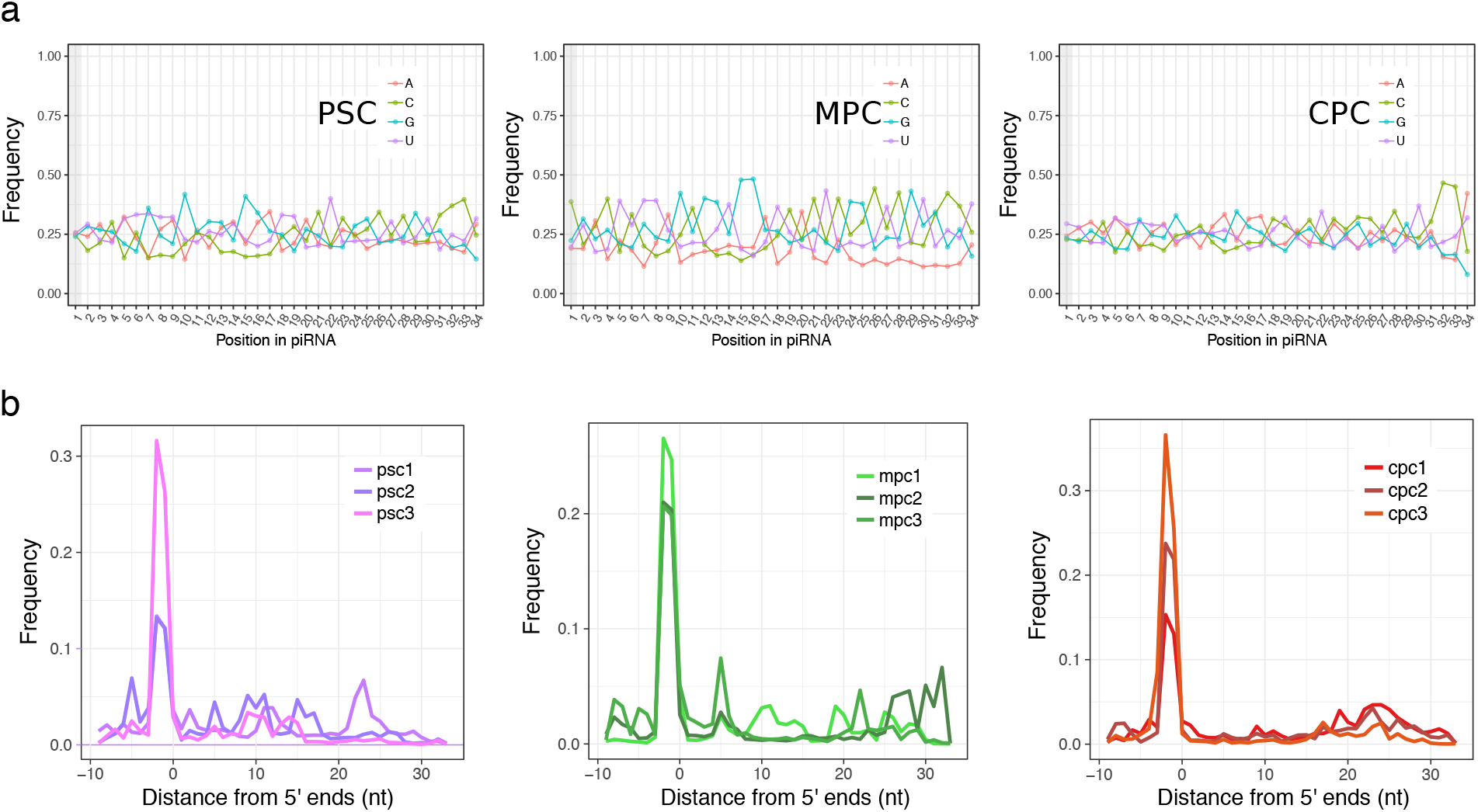
Characterization of reads mapped to known piRNAs. a) Frequency of bases per position in reads mapped to known piRNAs after fully processing alignments from PSC, MPC and CPC samples (merged replicates). Position 1 is marked by a vertical grey bar. b) Frequency profile showing overlap between reads mapped to known piRNA loci in sense orientation and complementary reads using an independent approach (ssviz package).

### Expression of piRNAs during cardiac differentiation

To study the expression profile of known piRNAs in differentiating PSC we kept those with an average count among replicates higher than or equal to 3. Normalization by library depth showed equivalent distribution of relative quantifications between samples (Supplemental Figure 3a), enabling confident identification of 447 piRNAs considering the three cell differentiation stages investigated (Supplemental Table 2). Despite some differences between replicates, each stage of cell differentiation was categorically defined by a specif piRNA expression profile (Supplemental Figure 3b) which was also reflected in Principal Component Analysis results (Supplemental Figure 3c). These identifying profiles preferentially aggregated PSC and MPC together indicating a greater resemblance between samples of these two cell populations than with CPC.

Of the 447 identified piRNAs, 241 were expressed in PSC, 218 in MPC and 171 in CPC (Supplemental Table ??). Differential expression analysis revealed only 30 genes with significant shifts in RNA levels (−1 > log2FC > 1; −log10p-value > 1.30) for the comparison between PSC and MPC, while 137 were differentially expressed (DE) between PSC and CPC and 153 between MPC and CPC (Figure 3a). Of the total 447 piRNAs, 204 were found to be DE with respect to CPC, 86 of which were shared by MPC and PSC (Supplemental Figure 3d and Supplemental Table 2). These results were consistent with correlation analysis that showed a higher Pearson coefficient for the MPC-PSC pair (R=0.5, p<2.2e-16) than for CPC-PSC (R=−0.0062, p=0.9) (Figure 3b), suggesting that PSC bear a greater resemblance to MPC than to CPC not only in the identity of expressed piRNAs, but in their abundance as well. Upregulated piRNAs accounted for 14% of total DE piRNAs in CPC (Figure 3c and Supplemental Table 2), far fewer than the downregulated piRNAs (Figure 3d and Supplemental Table 2). We validated several piRNA transcripts (piR-1919272, piR-2519215 and piR-97458) by qPCR in an independent set of samples from H9 pluripotent cells and 14 days after the onset of cardiac differentiation, corroborating our detection pipeline and subsequent DE analysis (Supplemental Figure 4).

**Figure 3:**
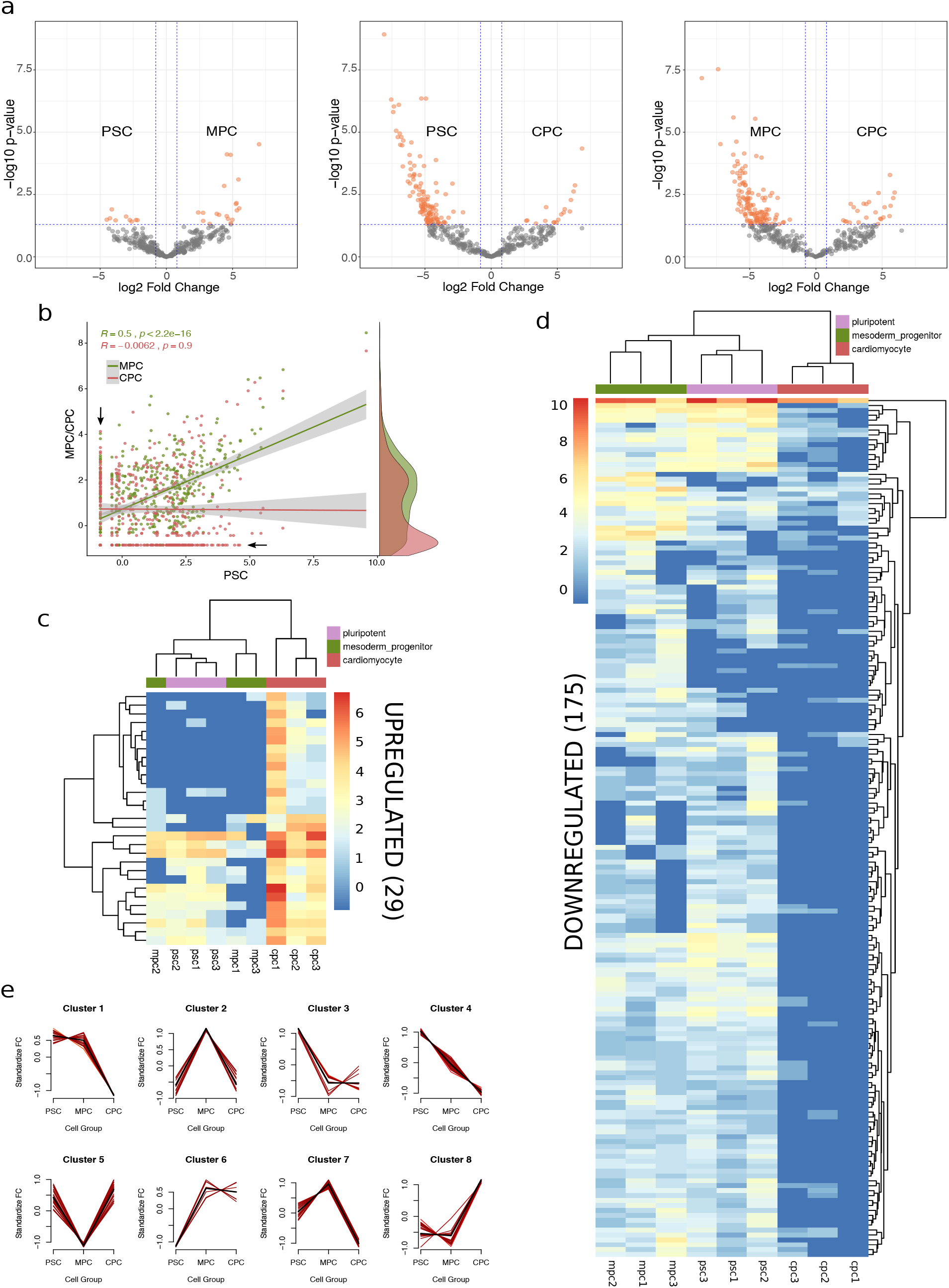
piRNA expression profile in differentiating pluripotent stem cells. a) Differential expression analysis performed on raw counts with DESeq2 package. Significantly different expression values (−1 >= log2FC >=1; −log10p-value > 1.30)) are represented as orange dots in the volcano plots for the three possible comparisons: MPC vs. PSC (left), CPC vs. PSC (center) and CPC vs. MPC (right). b) Scatter plot of normalized expression data showing Pearson correlation analysis on MPC (green dots) and CPC (red dots) versus PSC. Marginal density plots to the right denote areas of highly abundant data and black arrows mark the positions of the most DE piRNA genes. c) Heatmap shows normalized counts of upregulated piRNAs genes in CPC considering the three cell populations (PSC, MPC, CPC). Dendrograms resulted from running hard unsupervised clustering algorithms on piRNA genes (left) and samples (top). d) Heatmap as in c showing downregulated piRNA genes in CPC. e) Implementation of soft clustering algorithms (R package MFuzz) produces eight distinct patterns (clusters 1 to 8) of piRNA expression.

Differentially expressed piRNAs ranked among the top expressing piRNAs. This is probably due to the fact that highly expressed genes are inherently less sensitive to inter-replicate noise, hence more likely to return a lower p-value for contrasts. Thus, in order to investigate potential patterns underlying expression data which might have been masked from differential expression analysis, we implemented a soft clustering algorithm to data. This approach returned 8 different patterns of piRNA expression (Figure 3e and Supplemental Table 3), or Expression Clusters (EC), that reflected two dynamically relevant tendencies: down-regulation of piRNAs towards cardiac differentiation (cluster 1 to 4 and 7) and upregulation of piRNAs towards cardiac differentiation (cluster 5, 6 and 8). The former, as was previously observed, encompassed the majority of DE piRNAs. Regardless of the condition (up or downregulated) of piRNAs in CPC, it was clear that a fraction of piRNAs sustained early change (PSC to MPC) while others shifted later in the differentiation process (MPC to CPC). Interestingly, expression profile of human *PIWI* genes (*HIWI, HILI, HIWI2* and *HIWI3*) changed between day 0 and 14 of differentiation (Figure 4a). While no conclusive results were obtained for *HIWI* and *HIWI3, HILI* and *HIWI2* were upregulated towards day 14 suggesting that there might be a connection between these *PIWI* genes and cardiac piRNAs. Specific markers of pluripotency and cardiomyocytes were measure at these timepoints to corroborate cell identity (Figure 4b). In addition, analysis of H9 and H1 published RNA-seq data validated upregulation of *HIWI2* with cardiac differentiation (Supplemental Figure 5, (*32*)).

**Figure 4:**
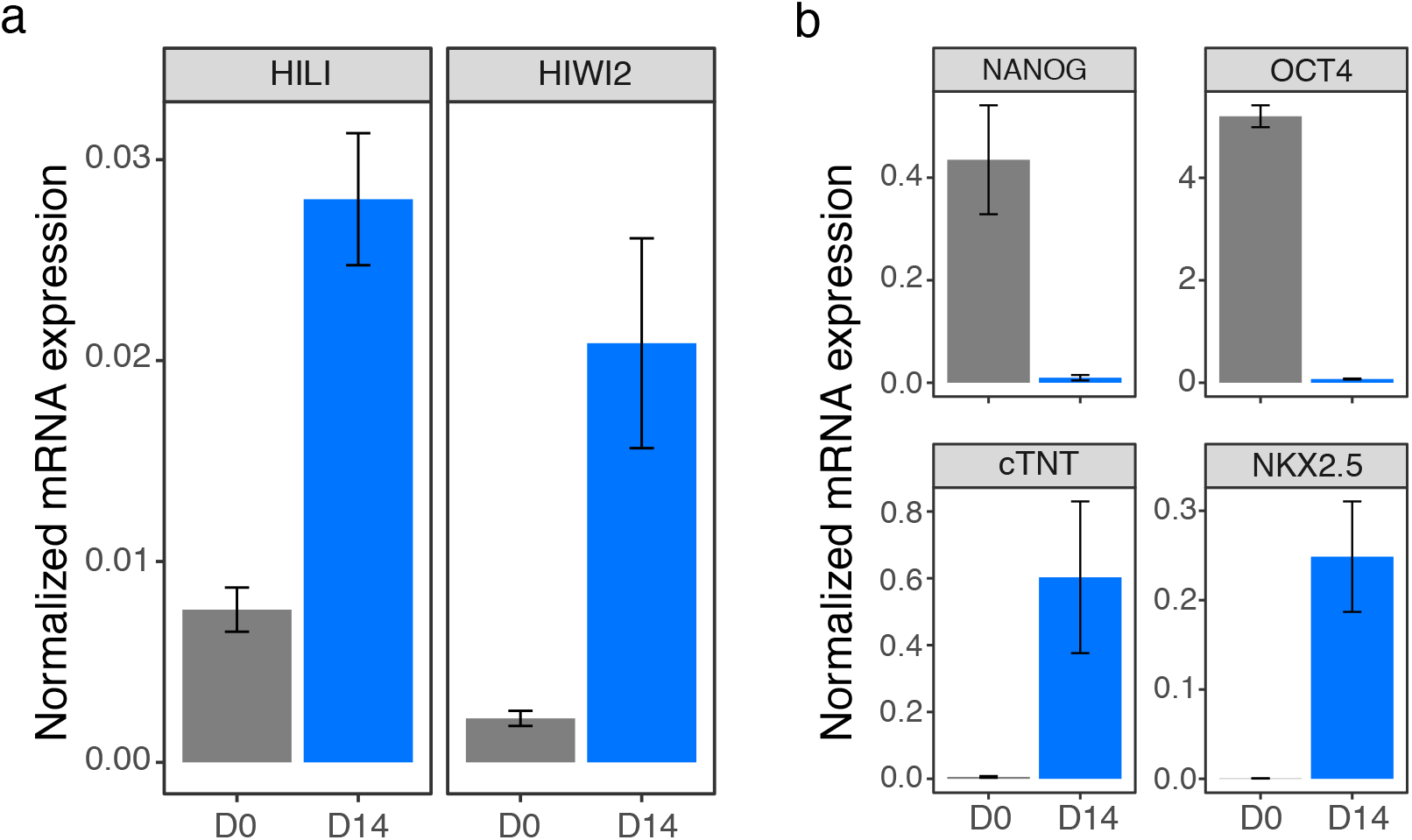
Expression of human *PIWI* genes in cardiac differentiation. a) Transcript levels of *HILI* (PIWIL2) and *HIWI2* (PIWIL4) were measure by qPCR in H9 pluripotent cells (D0) and H9-derived cardiac progenitors (D14). b) Stage-specific markers were evaluated by qPCR in samples from a. Pluripotency genes are shown in the top row and genes for cardiac lineage in the bottom. All results were expressed as mean±se of two independent experiments after normalization by the geometric mean of *RPL7* and *HPRT1* housekeeping genes.

### Genome distribution of expressed piRNAs

Identified piRNAs were distributed rather uniformly throughout the nuclear genome (Figure 5a), except in chromosome Y for which no data was available given the XX karyotype of H9 embryonic stem cells. Moreover, Expression Clusters did not seem to follow any particular arrangement in these chromosomes as well (Figure 5a, center of circular plot). Inclusion of the mitochondrial chromosome (chrM) in the analysis revealed that 90 of 447 piRNAs originated from the mitochondrial genome (Figure 5b). This was consistent with previous work in human somatic cancer cell lines reporting the synthesis of piRNAs from mitochondrial genome ((*33*)). In fact, the chrM was the major contributor of expressed piRNAs in our samples and was mostly dominated by three EC: a) cluster 3, with piRNAs highly expressed in PSC; b) cluster 8, with piRNAs highly expressed in CPC; c) cluster 5, with piRNAs highly expressed in both PSC and CPC.

**Figure 5:**
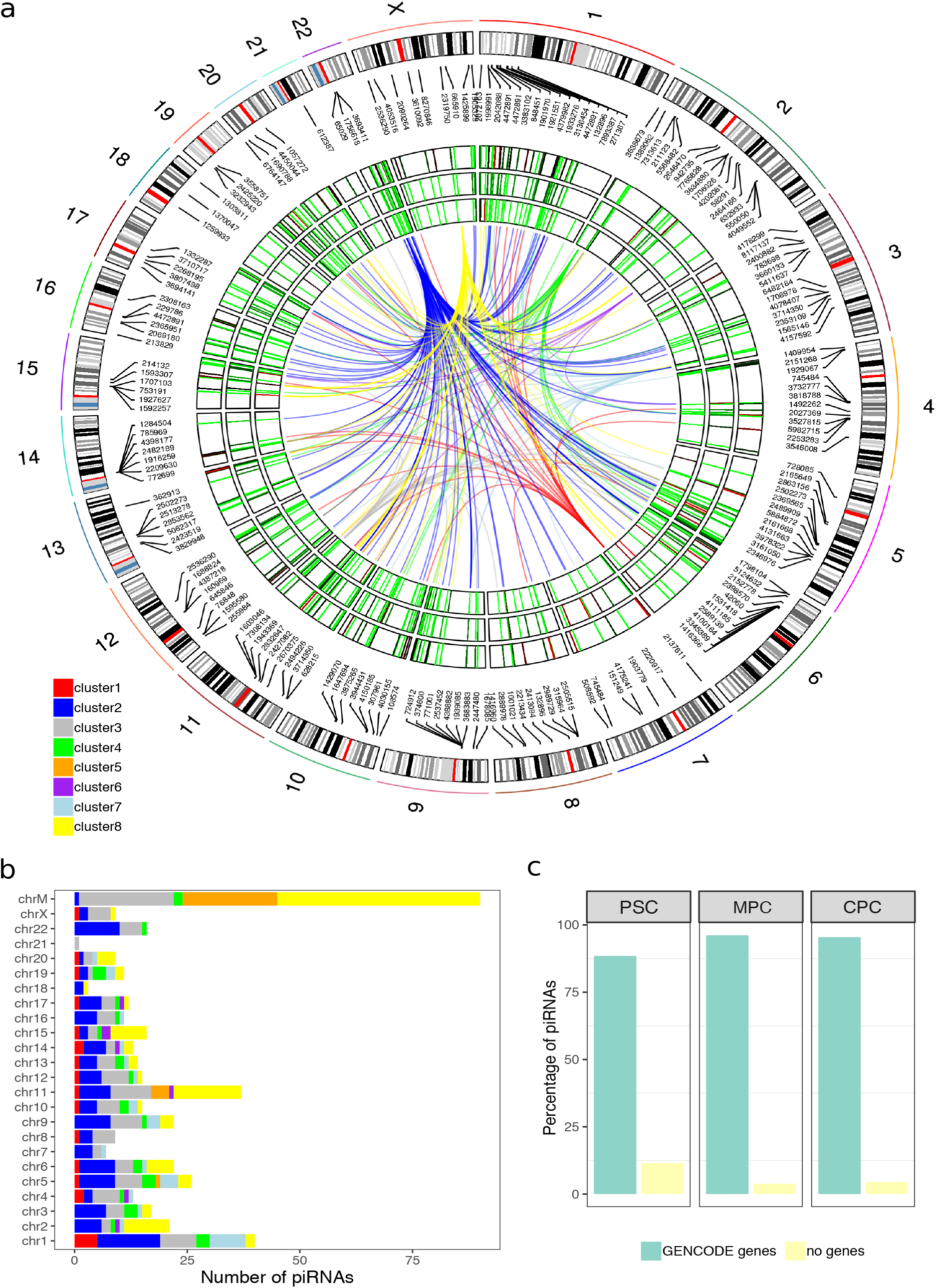
Distribution of piRNAs and Expresion Clusters in the genome. a) Normalized expression of piRNAs from averaged samples of PSC (outer track), MPC (middle track) and CPC (inner track) displayed lengthwise in a circular representation of human autosomal chromosomes (1 to 22) and chromosome X. Level of expression is depicted as heatmaps inside the tracks and follow a red-green scale for high expression and low expression respectively. The center of the plot shows links between piRNAs that belong to the same Expression Cluster (EC); color key for the links appears at the bottom left corner of the plot. b) Bars represent number of piRNAs expressed in all samples per chromosome. EC membership of piRNAs is indicated with color key as in a (top left corner of barplot). c) Percentage of piRNAs in PSC, MPC and CPC samples that intersected or not with genes from GENCODE database (v29).

We corroborated this finding by evaluating the distribution of mapped reads over chrM, and thus eliminated the possibility of errors during counting of reads per transcript (Supplemental Figure 6a). Despite our pipeline for identification of expressed piRNAs filtered out all reads that mapped to ncRNAs-other than piRNAs-using DASHR database, 90% of mitochondrial piRNAs (81 out of 90) mapped directly to tRNA and rRNA annotations (GENCODE v29) (Supplemental Figure 6b). DASHR database showed only one annotation in chrM (LSU-rRNA) that corresponded to the large ribosomal subunit RNR2 (Supplemental Figure 6c). This was not the case in the nuclear genome where no piRNAs were found to map on rRNAs and tRNAs annotated in GENCODE database (Supplemental Figure 6d). Nonetheless, piRNAs identified in length-filtered data (initial step of filtering, Figure 1b) did not map to nuclear rRNA or tRNA annotations from GENCODE to begin with (Supplemental Figure 6d), suggesting that this step was sufficient enough to remove reads mapping on them.

Regardless of the chromosome distribution, identified piRNAs localized preferentially on gene annotations (Figure 5c). PSC samples showed that 88.5% of piRNAs were generated from gene features, while the percentage was higher in MPC and CPC samples, with 96.2 and 95.5% respectively.

### Protein coding and lncRNA genes hosting piRNAs

Further analysis on genomic distribution of identified piRNAs revealed that nearly 65% of those intersected to gene features originated from coding (53%) and long non-coding (12%) annotations (Suplemental Figure 5d). To test whether these events were random, we shuffled our samples 1500 times (bootstraping) and analyzed intersection to these features in sense and antisense orientation. Once data was collected, we calculated enrichment on genes as “sample piRNAs” over “shuffled piRNAs” and determined that sense-oriented piRNAs occurred non-randomly on protein coding and long non-coding (lnc) genes (Figure 6a). On the contrary, intersection in antisense had poor fold enrichment values suggesting piRNAs were preferentially located elsewhere. We observed similar results for piRNAs identified in all three cell differentiation stages studied in this work, as well as in two samples (isogenic replicates) downloaded from ENCODE project corresponding to H1-derived neural progenitor cells (NPC). Both neural samples were handled following the same steps and criteria described before (Supplemental Figure 7).

**Figure 6:**
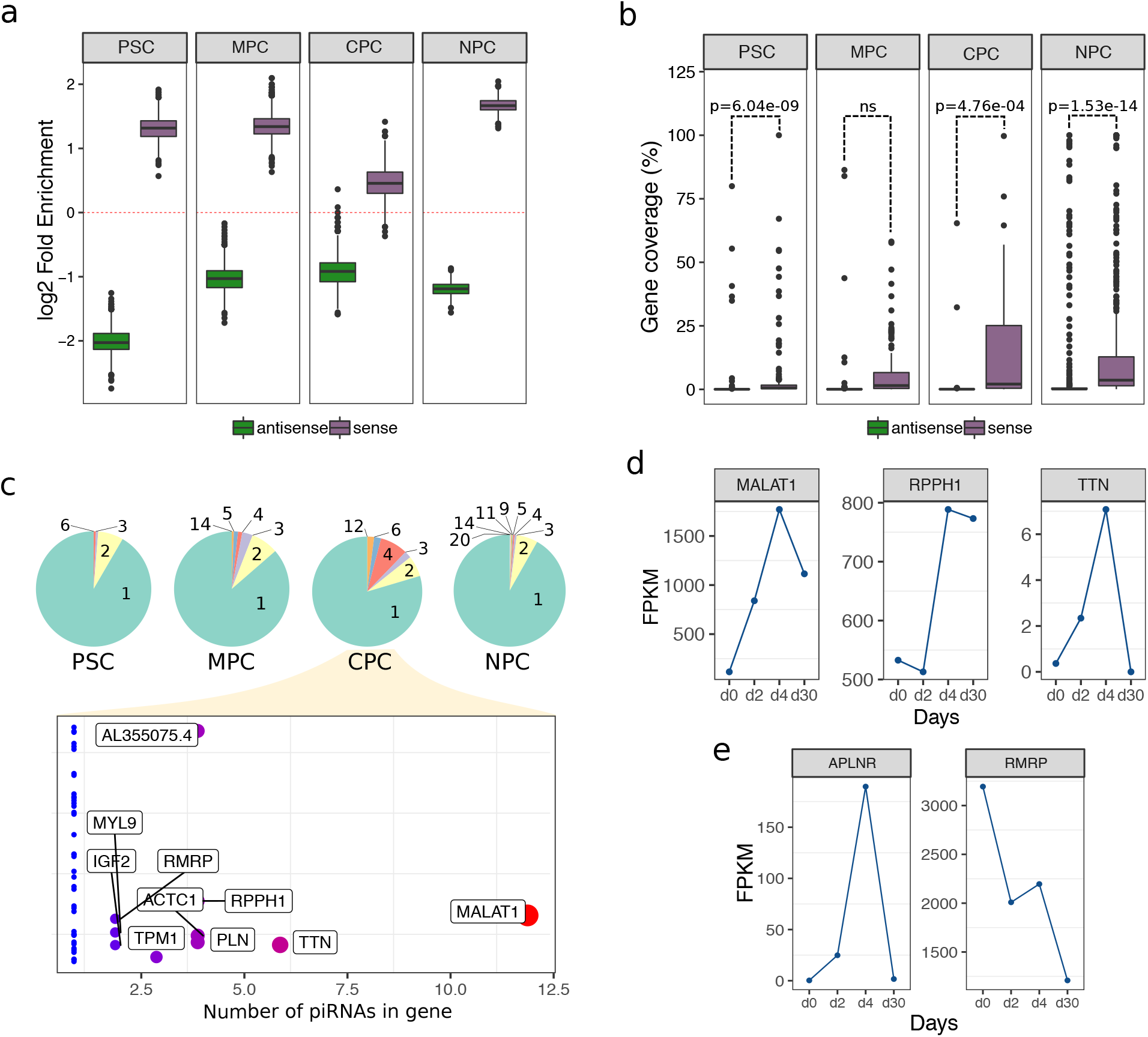
Proteing coding and lncRNA-originated piRNAs. a) Fold enrichment was calculated as number of piRNAs intersected to protein coding and lncRNA genes in sense (purple) and antisense (green) orientation over 1500 different random distributions (bootstraping). Red dashed line marks the point of no enrichment (log2 1=0). NPC: human neural progenitor cells. b) Distribution of percent coverage values on genes in sense (purple) and antisense orientation (green). P-value (p) for statistical analysis is shown in the plots (ANOVA). ns: not significant. c) Pie charts show the proportion of genes containing piRNAs. The number of sense piRNAs per gene in each category is indicated in the charts. Plot at the bottom provides detail on pie chart for CPC labeling genes with two or more piRNAs. Mitochondrial genes are not included in the analysis. d) Normalized counts (FPKM) for three genes from c during differentiation of H9 cells (day 0, day 2, day 4 and day 30) to CM. Data was extracted from previously published total RNA-seq experiments. e) Normalized counts (FPKM) for two genes containg piRNAs in MPC.

Taking into consideration that reads originated from protein coding and lncRNA features might have been the result of ordinary transcript degradation, we investigated the distribution of reads mapped on such piRNA-hosting genes. Results showed that the percentage of bases covered by sense-oriented reads in these genes was low, with a median value of 0.56% in PSC, 1.50% in MPC, 2.08% in CPC and 3.63% in NPC (Supplemental Figure 8a). Moreover, coverage was localized to a set of specific piRNAs instead of all piRbase annotations described in any single gene (Supplemental Figure 8b), proving to be inconsistent with random degradation-produced reads. Coverage by antisense-oriented reads was closer to none (Supplemental Figure 8c) and significantly lower than sense-oriented coverage in all cell population except in MPC (Figure 6b), possibly due to a higher level of dispersion in values of these samples.

The wide majority of piRNA-hosting protein coding and lncRNA genes harboured a single piRNA transcript with a tendency to augment the number of piRNAs per gene throughout the differentiation process (Figure 5 6c, pie charts). Like MPC and CPC, NPC exhibited a wider spectrum of piRNAs per gene than undifferentiated pluripotent cells (PSC). A more detailed exploration into CPC results revealed that *MALAT1* (lncRNA gene) and *TTN* (protein coding gene) contained the highest number of piRNAs −12 and 6 respectively-, followed by *PLN, RPPH1, ACTC1* and *AL355075.4* with 4 (Figure 5 6c, bottom panel). Using RNA-seq data from H9 cells differentiated to CM (*32*) we analyzed the expression profile of these piRNA-containing genes. Transcript abundance of *MALAT1, TTN* and *RPPH1* increased from day 0 (PSC) to day 2/4 (MPC) and then dropped between day 4 and day 30 (CPC), in consonance with the expression dynamics of piRNAs from EC 8 (Figure 6d). *PLN* and *ACTC1* RNA levels increased from day 4 to day 30 practically impervious to piRNA production, though lack of data between day 4 and 30 hindered our analysis for these genes (data not shown). With respect to *AL355075.4* gene, we found no count data available in RNA-seq samples. However, this gene overlaps *RPPH1* in sense orientation and it partially overlaps protein coding gene *PARP2* in antisense orientation, meaning that piRNAs originated from it could be potentially involved in regulating both genes. In fact, *PARP2* expression dynamic showed a steady descent in transcript levels from day 0 to 30, which is also consistent with the fact that the piRNAs originated from *AL355075.4* were also detected in MPC (Supplemental Figure 8d). Similar results were found when we studied two genes with high piRNA content (>3) in MPC (Figure 6e) -*APLNR* and *RMRP*-in which almost all piRNAs belong to EC 2. Taken together, these evidences suggest that piRNAs originated from these genes may be implicated in their downregulation or possibly in a moderate fine-tuning as in the case of *PLN* and *ACTC1*.

### Functional analysis on piRNA-hosting genes in differentiated cells

The expression profile of piRNAs proved to be sufficient to clearly discriminate CPC samples not only from PSC and MPC populations, but from neural progenitors (NPC) as well. The comparison between CPC and NPC samples revealed that 52 piRNAs were expressed in both types of differentiated cells, but more importantly the majority were not (Figure 7a). Unshared piRNAs constitute a unique repertoire for each cell population which could possibly reflect upon diverse functional processes. To evaluate this notion, we extracted all protein coding genes which were intersected by at least one piRNA and determined their involvement in any biological process (BP). In search for overrepresented terms (BPs with more genes involved than expected), we found that CPC and NPC showed markedly different terms. The BPs associated to CPC were intimately related to heart development and muscle differentiation and contraction (Figure 7b), while overrepresented BPs in NPC showed a clear inclination towards neurogenesis regulation and neural proliferation and development (Figure 7c). The group of genes intersected by piRNAs shared by both CPC and NPC (52 piRNAs in venn diagram) did not participate in any of the statistically significant overrepresented BPs, meaning that enriched categories for each population are mostly based in their unique collection of piRNAs.

**Figure 7:**
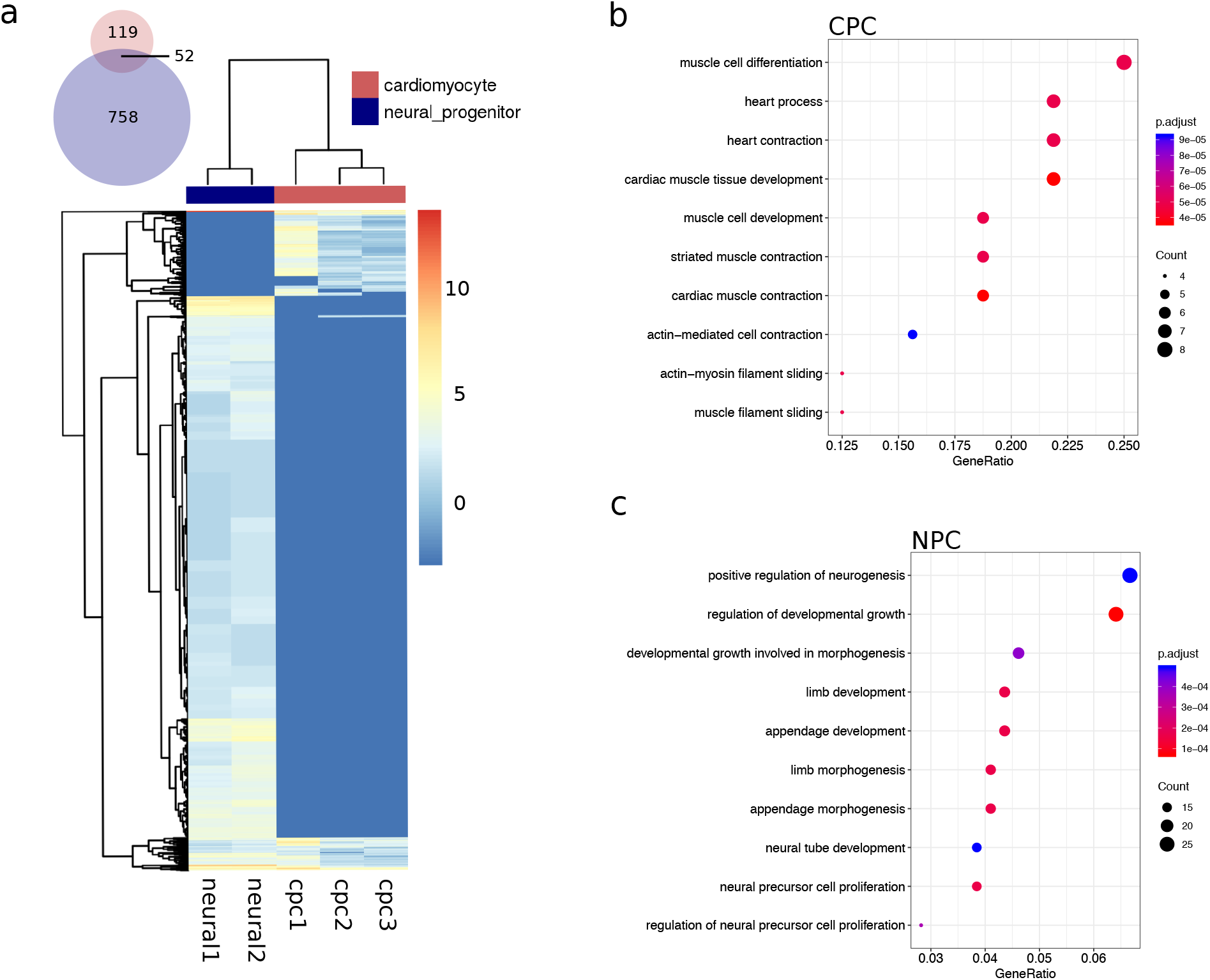
Functional exploration of piRNAs in differentiated cells. a) Expression profiles of piRNAs (log2 normalized counts) in CPC (cpc1, cpc2 and cpc3) and NPC (neural1 and neural2). Color key for sample clustering is displayed at the top right corner of the heatmap. b) Overrepresented biological processes (BPs) on proteing coding genes harbouring piRNAs from CPC samples. c) Overrepresented BPs on proteing coding genes harbouring piRNAs from NPC samples.

## Discussion

Since the first mechanistic evidences of piRNA biogenesis in ovarian follicle cells of *D. melanogaster* were published (*14*), much information has emerged on the somatic expression and function of this type of regulatory small RNAs. Several publications demonstrated that piRNAs (or piRNA-like RNAs) originate from discrete genomic regions of somatic cells in a wide diversity of species and tissues (*15, 27, 28, 34, 35*). In agreement with this line of evidence, we report the expression of 447 small RNAs consistent with piRNAs among three stages of differentiation of pluripotent stem cells to cardiomyocytes using a database-driven approach.

The pipeline leading to the identification and quantification of piRNAs involved two filtering steps that were implemented to avoid innacurate interpretation of results. Firstly, aligned reads shorter than 24nt and longer than 34nt were discarded from further analyses. However, our results showed a higher-than-basal frequency of 36-37nt-long reads, which prompts the issue if these reads should have been kept for further investigation as potential longer piRNAs or perhabs remnants of piRNA precursors. Length restriction answers to one of the hallmark attributes of mature piRNA transcripts, albeit the range seems to vary across species. In fact, in *C. elegans* 21nt-long piRNAs are produced from transcript precursors of 25-27nt in length (*36*) whose processing machinery is partially unknown. The reason for the range diversity has been convincingly connected to the activity of the proteins involved in their biosynthetic pathway (biogenesis). It is possible that they also contribute to explaining the differences between germline and somatic piRNAs considering that diverse sets of enzimes have been reported to be engaged in piRNA synthesis in these cell lineages (*36, 37*). Moreover, the differential expression profile of *HIWI* genes observed in our data -and in H9 and H1 external RNA-seq data-points to cell-type specific functions for these proteins and consequently for their piRNA partners.

In a second step, length-filtered reads mapping to small ncRNAs other than piRNAs were removed from samples. The fact that remaining reads in PSC and CPC samples showed a moderate 5’ U bias, whereas MPC samples did not could probably be related to the transitional nature of this cell population. However, none of the samples exhibited the characteristic 10nt-overlap signature of secondary piRNAs, which is a generally accepted feature in germline cells of most animals where piRNAs are synthesized through both primary and secondary (ping-pong loop) pathways (*15, 26*). Despite synthesis in somatic cells has been proposed to produce only primary piRNAs, many of the mechanisms underlying piRNA biogenesis-specially in non-gonadal tisues-are not yet fully understood. Conceivably, the filtering steps eliminated potential piRNAs from the three stages, though it has been argued that a considerable amount of annotated piRNAs are actually ncRNA fragments derived from rRNAs, tRNAs and even miRNAs (*31*). On the matter, an insightful disscussion by Tosar and collaborators (*31*), advocating for gonadal piRNAs, suggested that somatic piRNAs mapping to ncRNA fragments are not unquestionably wrong, still further biochemical evidence is needed to include them as such.

An important aspect of this work lies on the identification of a piRNA expression profile associated to each of the cell populations under study. These expression profiles parallel the embryological connection between the stages, where PSC is more closely related to MPC than to CPC (*2, 38*). Upon this premise, the piRNAs identified as early-changing could potentially be involved in maintaining pluripotency or in the commitment of pluripotent cells to mesoderm progenitors, which might eventually differentiate to multiple lineages. Late-changing piRNAs, on the contrary, would influence further commitment of mesoderm cells to cardiac progenitors. The fact that six times more piRNAs were downregulated rather than upregulated during differentiation to CPC suggests that piRNA pathways become less relevant in differentiated cells. It is possible that mechanisms evolutionarily linked to the regulation of transposable elements are not as critically conserved in differentiated cells as they do in cells with high proliferation rates or reproductive functions, such as pluripotent cells and germ line cells. For example, it has been proposed that cancer cells might promote piRNA biosynthetic pathways as a mechanism to reduce genome instability caused by increased mitotic and transcriptional activities (*39*).

The genomic localization of piRNAs included in anyone particular EC was not the same. In fact, piRNA expression appeared to be uniformly scattered across the genome except for the mitochondrial chromosome. The majority of piRNAs identified in the mitochondrial genome mapped to rRNAs or tRNAs and though we have not definitively proved they are truly piRNAs, previous work established a link between tRNA-/rRNA-derived piRNAs, *HIWI2* expression and regulation of metabolic processes in somatic cells (*40*). Analogously, the increased levels of piRNAs from mitochondrial tRNAs/rRNAs and the significant upregulation of *HIWI2* (day 14 v. day 0, and external H9 RNA-seq data) in CPC could seemingly be connected to the large-scale modifications in CM metabolism (*41*).

Even though identified piRNAs were dispersed throughout the genome, it was clear that the vast majority of them originated from gene loci. However, it is not yet clear the reason why these piRNAs are generated from the sense strand of their hosting genes. One possibility relies on the capacity of PIWI/piRNA complexes to direct recruitment of DNA and histone methyltransferases, modifying accesibility of transcriptional machinery to chromatin (*25, 42*). Available data of DNA or histone methylation status in the three stages of cardiac differentiation is scarse or dissimilar, so preliminary correlation analysis were not conclusive at this point (data not shown). Nevertheless, further experiments on promoter methylation and H3K9me3 mark deposition ought to be performed to pursuit this possibility. Also, considering that antisense transcripts have been described to positively regulate stability of sense RNA (*43*), it is possible that sense-originated piRNAs regulate antisense transcript levels in a miRNA-like mechanism. For instance, *TALAM1*-an antisense transcript at the *MALAT1* gene locus-promotes stability and maturation of Malat1 RNA by facilitating enzymatic cleavage of its 3’ end (*43*), thus a potential piRNA-mediated downregulation of *TALAM1* would redound to diminished *MALAT1* levels.

In sum, the evidences presented here contribute to understanding the dynamic expression of piRNAs during differentiation of pluripotent stem cells to cardiomyocytes and further explore their potential function as post-transcriptional modulators in somatic cells. Together with miRNAs, piRNAs seem to participate in the fine-tuning of transcript levels, adding yet another layer to the complex and intrincated networks governing gene expression.

## Methods

### Small RNAseq data

Data samples used in this work (PSC: H9 human embryionic stem cells, MPC: early mesoderm progenitor and CPC:cardiomyocytes) were generated in our laboratory following previously described protocols and are available under accession code GSE108021. Briefly, H9 cells (H9-hTnnTZ-pGZ-D2 obtained from WiCell) were routinely maintained in co-culture with irradiated primary mouse embryonic fibroblasts. Mesoderm induction (MPC population) was performed by initially seeding cells with mTeSR (StemCells Technologies) on plates coated with Geltrex (Thermo Fisher Scientific) and then switching to (*2*) StemPro®-34 SFM (Thermo Fisher Scientific) supplemented with Activin A only at the first day, BMP4, VEGF and bFGF (Thermo Fisher Scientific) for 3.5 days. At this point, mesoderm progenitors were isolated by FACS with anti-CD326 and anti-CD56 (Biolegend). CPC population was obtained by formation of embryoid bodies with H9 cells using BMP4, bFGF and Activin A in StemPro®-34 followed by addition of VEGF and Wnt inhibitor, IWR-1. Libraries for small RNA sequencing were prepared with 200 ng of RNA using NEBNext Small RNA Library Prep Set with modified adaptors and primers compatible for Illumina (New England Biolabs). Single end sequencing was carried out at the TCGB Resources (UCLA Path and Lab Med) using an Illumina HiSeq 2500. Culture conditions and sequencing of small RNAs for these samples are more extensively described in (*11*).

### External data

Human testis small RNA-seq samples from two men of 54 and 37 years old (GSE88414 and GSE88124, respectively) and H1-derived neural progenitor cells (GSM1296459 and GSM1296460) were downloaded from ENCODE (encodeproject.org). RNA-seq data (counts per transcript) from H1 and H9 cardiac differentiation protocols can be found under GEO accesion GSE85331.

### Data processing and analyses

Adapters were removed from raw sequencing reads with cutadapt (v1.9.1) keeping reads with a minimum of 20 and up to 50 nt in length. Quality checked (FastQC) processed reads were mapped to human reference genome (GRCh38/hg38) using STAR aligner (v2.5.3a (*44*)) under mostly default parameters. Mapped reads in output SAM/BAM files were filtered by read length (23 < RL < 35) with samtools and custom awk scripting. Resulting reads were intersected (bedtools v2.27.1 (*45*)) to ncRNAs in a strand specific manner (DASHR (*46*)) to remove potential misleading alignments. Raw counts on piRNAs were determined with htseq-count matching mapped reads to piRNA coordinates downloaded from piRBase (*47*). Counts were then fed into DESeq2 for differential expression analysis (p<0.05 and fdr<0.1). Soft clustering methods were implemented with R package Mfuzz (v2.42.0 (*48*)) using parameter m=1.15. In parallel to our customized pipeline, tools for ping-pong signature detection like ssviz R package and PingPongPro (*49*) were run following recommendations from authors. Graphics and statistical analyses were performed in R software and deepTools (*50*).

### Reverse Transcription of piRNA

To obtain cDNA from piRNA transcripts we adapted a previously described methodology employed for miRNA detection and amplification (*51*). Briefly, stem-loop retrotranscription (RT) primers were generated using 6-8nt from the 3’ end of every piRNA of interest. Each RT reaction was performed with a maximum of 10 different stem-loop primers including one for RNU6B and hsa-miR-302b as controls. SuperScriptIII retrotranscriptase (Thermo Fisher Scientific) was used for RT reactions following guidelines from manufacturer. Detection by qPCR involved forward primers matching the sequence of target piRNAs and a reverse universal primer complementary to the stem-loop RT primer.

### Real time PCR

Total RNA was prepared with TRI-Reagent (Sigma Aldrich) following manufacturer’s instructions and then reverse transcribed using MMLV reverse transcriptase (Promega) and random primers for detection of polyadenilated transcripts. Quantitative real time PCR (qPCR) was performed in a StepOne Real Time PCR system (Applied Biosystems). Expression was normalized to the geometrical mean of HPRT1 and RPL7 housekeeping genes and log2 transformed. Statistical significance for qPCR results was analyzed by ANOVA followed by Tukey’s multiple comparison test. Primers sequences are available on request.

## Acknowledgments

We would like to thank Fundacion FLENI and Fundacion Perez Companc for their continuous support.

## Author Contributions

ALG and SGM design experiments and wrote manuscript. MAS, MCHC, NP, SC, CC, AMM, NLSV and CA performed global bioinformatic analysis. GN, XG, AW, LNM, GS, CL and SM discussed and reviewed analyses and manuscript. GS and SM provided fundings for this paper.

