## Supplementary FIles for "Analysis of the expression of PIWI-interacting RNAs during cardiac differentiation of human pluripotent stem cells"

**Alejandro La Greca**

### Supplementary Files

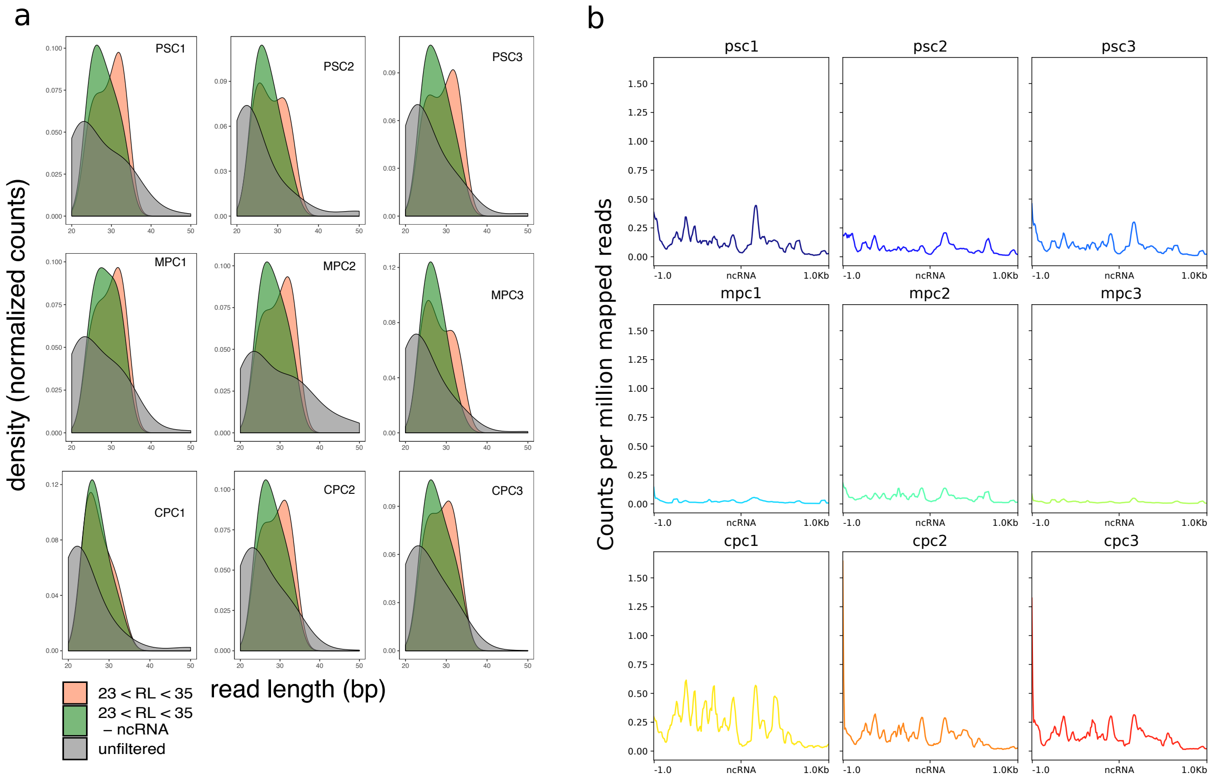


FIGURE S1: **Read length control in filtered samples and mapping to other non coding RNAs.** a) Read length expressed as a function of density for pluripotent (PSC1, PSC2 and PSC3), mesoderm progenitor (MPC1, MPC2 and MPC3) and cardiomyocytes (CPC1, CPC2 and CPC3) samples before filtering (unfiltered) and after filtering (23 < RL < 35 and -ncRNA). Color key is indicated in the plot. b) Analysis of coverage on non coding RNAs loci from DASHR database for fully processed normalized (counts per million) samples.

| sample | unfiltered (1) | length filtered (2) | length+ncRNA filtered (3) | 2/1 (%) | 3/1 (%) |
|---|---|---|---|---|---|
| **psc1** | 13,475,096 | 5,622,650 | 1,272,076 | 41.73 | 9.44 |
| **psc2** | 8,979,026 | 2,481,733 | 656,781 | 27.64 | 7.31 |
| **psc3** | 9,306,433 | 3,418,745 | 844,994 | 36.742 | 9.08 |
| **mpc1** | 18,690,310 | 8,922,435 | 3,494,534 | 47.74 | 18.705 |
| **mpc2** | 13,137,228 | 5,017,796 | 1,090,201 | 38.20 | 8.30 |
| **mpc3** | 14,426,323 | 5,582,968 | 1,921,400 | 38.70 | 13.32 |
| **cpc1** | 10,550,374 | 3,362,057 | 635,333 | 31.87 | 6.02 |
| **cpc2** | 10,863,324 | 4,833,395 | 882,061 | 44.49 | 8.12 |
| **cpc3** | 8,045,810 | 3,770,766 | 969,896 | 46.87 | 12.05 |

Table S1: **Number of mapped reads.** Reads were counted before processing (1) and after being filtered by length (2) and other ncRNAs (3). Remaining reads after processing are expressed as percentage(%) of unfiltered reads (2/1 and 3/1).

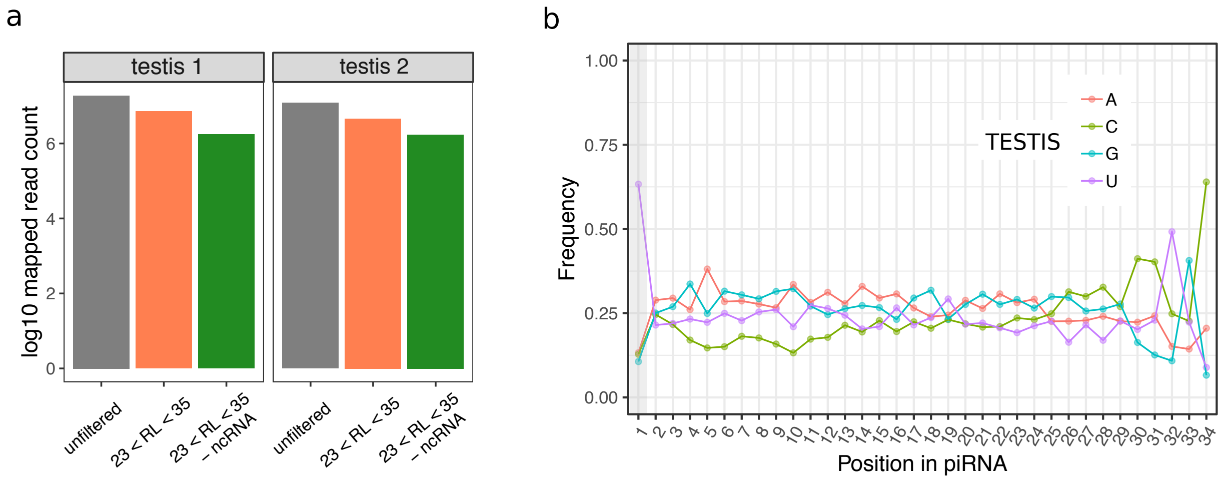


Figure S2: **Processing of testis samples.**. a) Number of mapped reads after employing the pipeline described in Figure 1 in human testis samples downloaded from ENCODE (merged replicates). b) Frequency of bases per position in processed mapped reads.

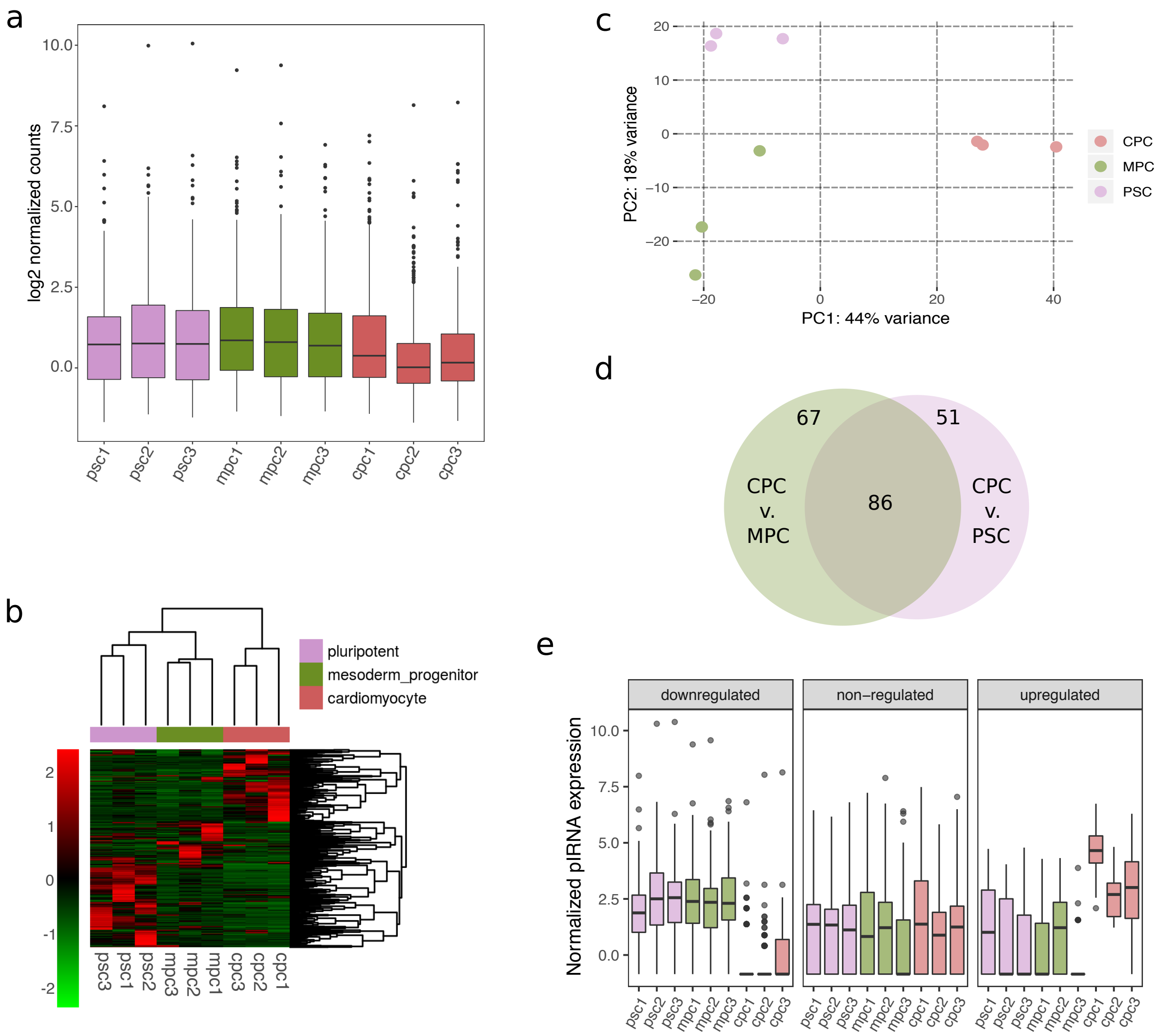


FIGURE S3: **Analysis of expression data and DE results.** a) Boxplot showing reads for the nine samples normalized by library depth and expressed as log2 counts per million (CPM). b) Heatmap in log2 CPM of piRNAs from a. Hard unsupervised clustering was performed on rows (piRNA ID) and columns (sample ID), and is shown as dendrograms. Color keys for heatmap and phenotype are indicated to left and in the top right corner of the graph, respectively. c) Principal Component Analysis performed on DESeq2 normalized counts. The color key is indicated to the right of the plot. d) Overlap of differentially expressed piRNAs in CPC versus MPC (153; green circle) and PSC (137; purple circle). e) Normalized expression of piRNAs upregulated, dowregulated and non-regulated with respect to CPC.

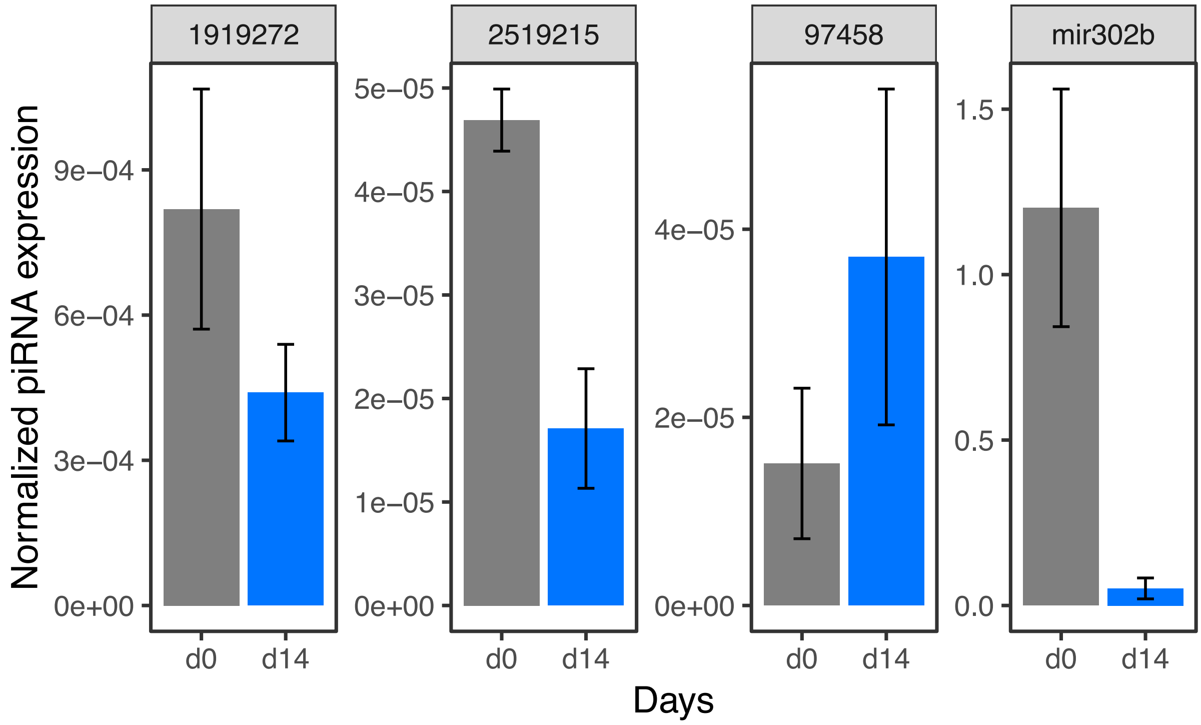


Figure S4: **Validation of piRNA expression.** Three piRNA transcripts (piR-1919272, piR-2519215 and piR-97458) were evaluated by qPCR using a specific retrotranscription protocol designed for small RNAs in day 0 and 14 of cardiac differentiation. Expression of mir302b -marker of pluripotency- was analyzed to assess protocol success.

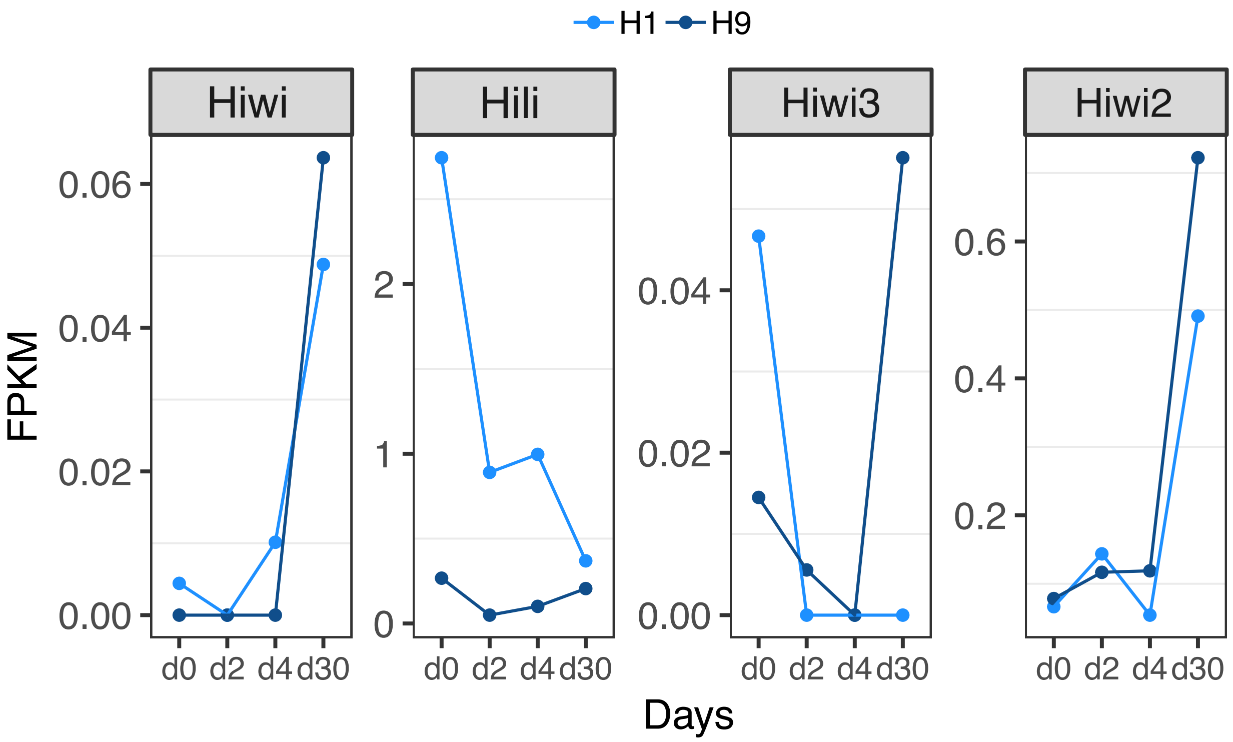


Figure S5: **Human Piwi genes expression profile in H1 and H9 embryonic stem cell lines.** Normalized RNA-seq counts (FPKM) from H1 and H9 cell lines were downloaded from GEO.

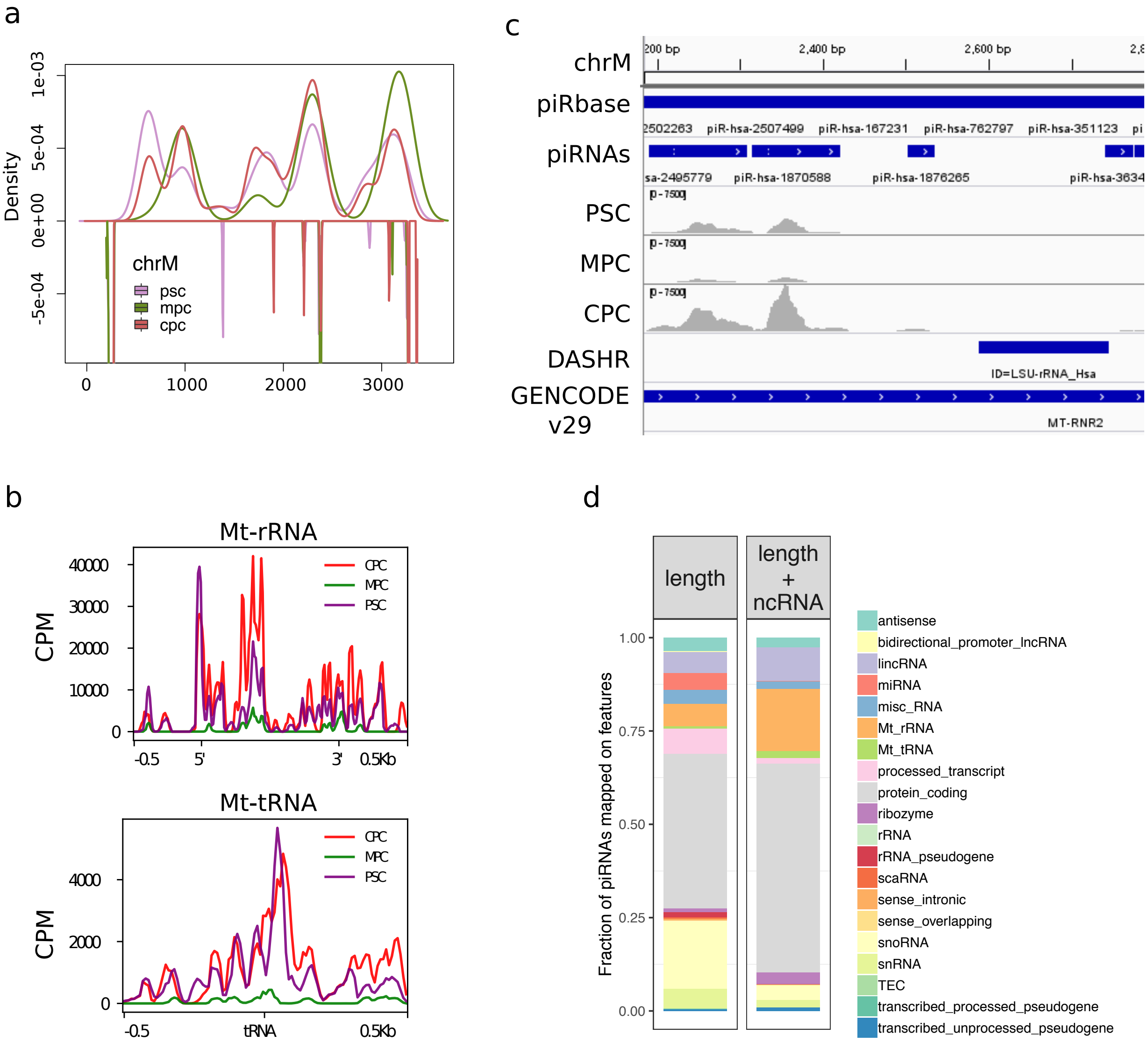


FIGURE S6: **Expression of piRNAs in mitochondrial chromosome.** a) Distribution of PSC, MPC and CPC mapped reads (merged replicates) as a function of density over a fraction of mitochondrial chromosome (chrM:1-4000). Profiles above zero correspond to plus strand and below zero to the minus strand. Color key is located at the bottom left corner of the plot. b) Coverage profiles in counts per million mapped reads (CPM) on the entire mitochondrial rRNA extension (MT-rRNA) and center of tRNA (MT-tRNA). Direction of rRNA genes are indicated by 5‘ and 3‘. c) Image captured from IGV software over a portion of the human mitochondrial chromosome (chrM:2,184-2,780). The tracks from top to bottom are: piRbase annotated piRNAs (piRbase), piRNAs identified in our samples (piRNAs), coverage profiles of PSC, MPC and CPC, DASHR database ncRNA annotations (DASHR) and GENCODE v29 gene annotations (GENCODE v29). d) Fraction of piRNAs mapped to genomic features annotated in GENCODE v29 database in length-filtered samples (length) compared to length+ncRNA-filtered (length+ncRNA) samples. Color key for features is indicated to the rigth of the bars.

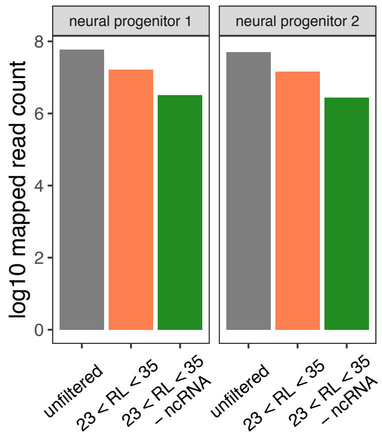


Figure S7: **NPC sample processing.** Number of mapped reads after employing the pipeline described in Figure **??** in neural progenitor samples downloaded from ENCODE project.

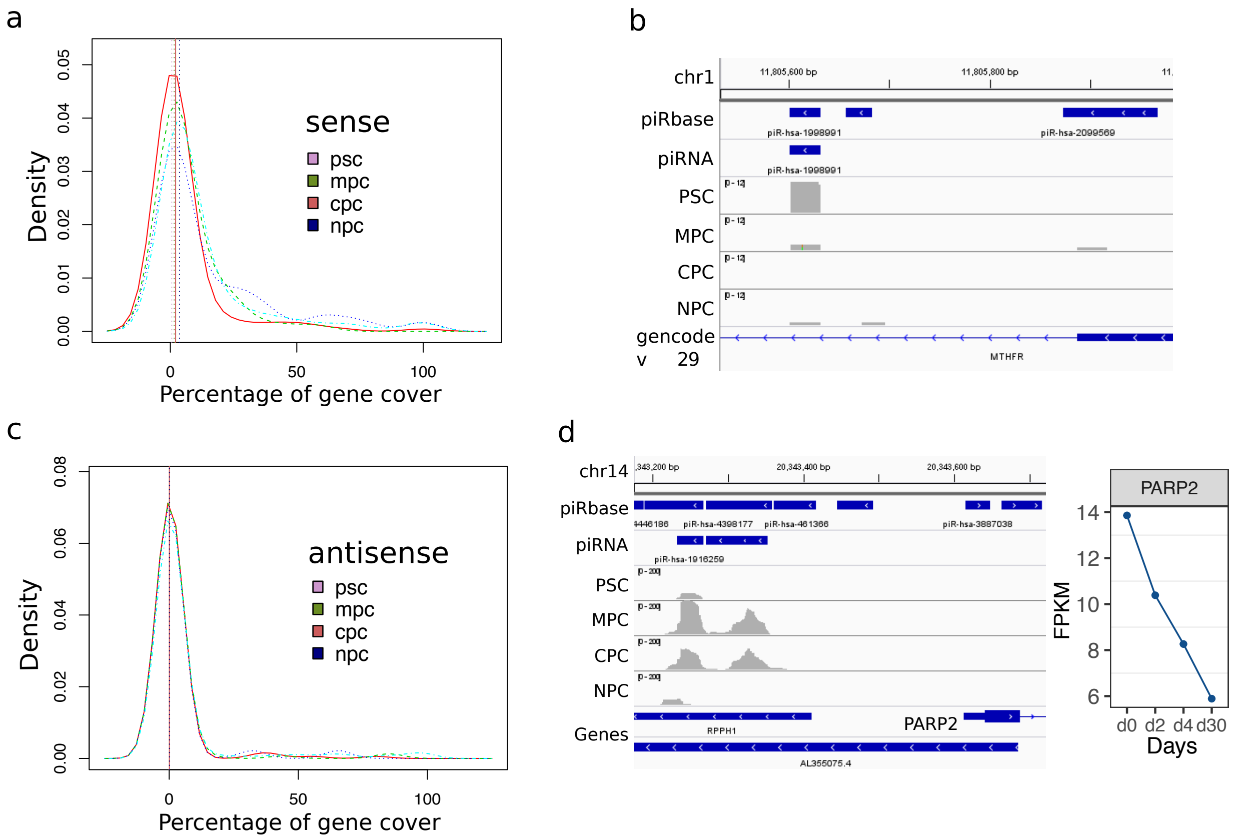


Figure S8: **Coverage on protein coding and lncRNA genes.** a) Desity estimation of percent (%) coverage on protein coding genes intersected by piRNAs in sense orientation. Vertical lines indicate medians of each curve. b) Image captured from IGV software portraying mapped reads of PSC, MPC, CPC and NPC samples on identified piRNAs. Tracks for piRNA annotation database (piRbase) and gene features (GENCODE v29) are shown. c) Density estimation as in a, in antisense orientation. d) Left panel shows IGV capture depicting piRNAs in PARP2 vecinity. Expression dynamic of PARP2 gene in RNA-seq samples from H9 cells differentiated to CM is shown to the right.

| | piRNA | psc1 | psc2 | psc3 | mpc1 | mpc2 | mpc3 | cpc1 | cpc2 | cpc3 |
|---|---|---|---|---|---|---|---|---|---|---|
| 1 | piR-hsa-1389062 | 252.26 | 1263.26 | 1335.60 | 666.72 | 757.88 | 83.67 | 110.68 | 260.73 | 280.30 |
| 2 | piR-hsa-4381848_2 | 86.38 | 70.78 | 77.08 | 78.63 | 235.99 | 77.84 | 81.81 | 31.18 | 79.49 |
| 3 | piR-hsa-3732777 | 17.49 | 35.39 | 110.80 | 105.40 | 106.53 | 60.32 | 24.06 | 55.58 | 73.21 |
| 4 | piR-hsa-1904126 | 12.72 | 40.01 | 49.38 | 96.20 | 72.82 | 83.67 | 57.75 | 39.76 | 89.25 |
| 5 | piR-hsa-1919272 | 50.88 | 26.16 | 33.72 | 10.87 | 72.82 | 3.89 | 178.05 | 41.12 | 60.66 |
| 6 | piR-hsa-3660133 | 88.50 | 67.70 | 77.08 | 44.34 | 64.73 | 35.03 | 1.60 | 0.00 | 0.00 |
| 7 | piR-hsa-3817390 | 25.97 | 69.24 | 49.38 | 23.42 | 31.02 | 60.32 | 0.00 | 1.36 | 2.79 |
| 8 | piR-hsa-151466 | 25.44 | 15.39 | 26.50 | 18.40 | 13.49 | 0.00 | 44.91 | 16.27 | 77.40 |
| 9 | piR-hsa-2539762 | 20.14 | 9.23 | 21.68 | 2.51 | 18.88 | 1.95 | 81.81 | 22.14 | 41.14 |
| 10 | piR-hsa-4091280 | 22.79 | 112.32 | 28.90 | 5.02 | 6.74 | 25.30 | 0.00 | 0.00 | 0.70 |
| 11 | piR-hsa-1939085 | 0.00 | 0.00 | 0.00 | 148.90 | 2.70 | 3.89 | 33.69 | 6.78 | 2.09 |
| 12 | piR-hsa-169217 | 49.29 | 26.16 | 56.60 | 19.24 | 29.67 | 9.73 | 0.00 | 0.90 | 2.09 |
| 13 | piR-hsa-97458 | 0.00 | 3.08 | 3.61 | 22.59 | 57.99 | 93.40 | 0.00 | 0.00 | 0.00 |
| 14 | piR-hsa-665910 | 1.06 | 0.00 | 1.20 | 2.51 | 0.00 | 7.78 | 14.44 | 14.01 | 131.08 |
| 15 | piR-hsa-2346976 | 19.61 | 60.01 | 30.11 | 22.59 | 22.93 | 9.73 | 0.00 | 0.00 | 0.70 |
| 16 | piR-hsa-1933276 | 2.65 | 0.00 | 3.61 | 107.08 | 36.41 | 13.62 | 0.00 | 0.45 | 1.39 |
| 17 | piR-hsa-3634065 | 5.30 | 6.15 | 8.43 | 0.00 | 10.79 | 0.00 | 105.87 | 6.78 | 19.52 |
| 18 | piR-hsa-2489909 | 0.00 | 3.08 | 3.61 | 6.69 | 5.39 | 114.81 | 4.81 | 3.61 | 7.67 |
| 19 | piR-hsa-3658275 | 14.84 | 7.69 | 3.61 | 1.67 | 16.18 | 3.89 | 67.37 | 14.91 | 18.83 |
| 20 | piR-hsa-2832647 | 7.42 | 46.16 | 6.02 | 7.53 | 2.70 | 27.24 | 20.85 | 14.91 | 5.58 |
| 21 | piR-hsa-3352181 | 12.72 | 16.93 | 10.84 | 89.51 | 2.70 | 1.95 | 0.00 | 0.00 | 2.09 |
| 22 | piR-hsa-151249 | 7.95 | 10.77 | 19.27 | 40.15 | 16.18 | 15.57 | 1.60 | 18.98 | 4.18 |
| 23 | piR-hsa-316012 | 4.77 | 13.85 | 6.02 | 2.51 | 10.79 | 31.13 | 20.85 | 14.46 | 26.50 |
| 24 | piR-hsa-2479371 | 21.73 | 10.77 | 14.45 | 5.02 | 24.27 | 1.95 | 30.48 | 8.59 | 10.46 |
| 25 | piR-hsa-611204 | 6.89 | 0.00 | 8.43 | 51.87 | 45.85 | 9.73 | 1.60 | 0.00 | 1.39 |
| 26 | piR-hsa-2513278 | 10.07 | 9.23 | 19.27 | 27.61 | 39.11 | 13.62 | 3.21 | 0.00 | 1.39 |
| 27 | piR-hsa-3974794 | 32.86 | 18.46 | 21.68 | 15.89 | 6.74 | 15.57 | 3.21 | 1.81 | 4.88 |
| 28 | piR-hsa-2780538 | 15.37 | 35.39 | 24.09 | 7.53 | 9.44 | 25.30 | 0.00 | 0.90 | 2.79 |
| 29 | piR-hsa-1872085_4 | 4.24 | 26.16 | 8.43 | 25.10 | 4.05 | 31.13 | 3.21 | 5.87 | 8.37 |
| 30 | piR-hsa-2526525 | 6.36 | 4.62 | 6.02 | 0.00 | 5.39 | 0.00 | 73.79 | 2.26 | 11.16 |
| 31 | piR-hsa-745484 | 2.12 | 3.08 | 0.00 | 62.74 | 2.70 | 0.00 | 33.69 | 3.61 | 0.70 |
| 32 | piR-hsa-147461 | 5.83 | 3.08 | 6.02 | 32.62 | 18.88 | 29.19 | 1.60 | 7.68 | 0.70 |
| 33 | piR-hsa-4460706 | 11.66 | 4.62 | 1.20 | 0.00 | 10.79 | 1.95 | 41.71 | 15.36 | 16.73 |
| 34 | piR-hsa-1927965 | 11.13 | 12.31 | 3.61 | 0.00 | 10.79 | 0.00 | 51.33 | 2.26 | 9.76 |
| 35 | piR-hsa-4303719 | 13.78 | 24.62 | 36.13 | 14.22 | 2.70 | 9.73 | 0.00 | 0.00 | 0.00 |
| 36 | piR-hsa-114666 | 18.02 | 15.39 | 26.50 | 1.67 | 9.44 | 29.19 | 0.00 | 0.45 | 0.00 |
| 37 | piR-hsa-1870588 | 18.02 | 1.54 | 9.63 | 1.67 | 16.18 | 0.00 | 35.29 | 9.49 | 8.37 |
| 38 | piR-hsa-7760463 | 1.59 | 0.00 | 0.00 | 33.46 | 55.29 | 3.89 | 1.60 | 0.00 | 4.18 |
| 39 | piR-hsa-1706026 | 1.06 | 4.62 | 0.00 | 74.45 | 9.44 | 0.00 | 8.02 | 0.00 | 1.39 |
| 40 | piR-hsa-2856604 | 4.24 | 23.08 | 24.09 | 25.93 | 9.44 | 7.78 | 0.00 | 0.45 | 2.09 |
| 41 | piR-hsa-1233052 | 17.49 | 3.08 | 7.23 | 0.00 | 12.14 | 0.00 | 41.71 | 9.04 | 4.18 |
| 42 | piR-hsa-2482189 | 2.12 | 10.77 | 0.00 | 49.36 | 6.74 | 3.89 | 14.44 | 3.16 | 3.49 |
| 43 | piR-hsa-2519215 | 15.90 | 18.46 | 37.33 | 7.53 | 5.39 | 7.78 | 0.00 | 0.00 | 0.00 |
| 44 | piR-hsa-2863156 | 5.30 | 15.39 | 6.02 | 20.08 | 4.05 | 35.03 | 4.81 | 0.90 | 0.70 |
| 45 | piR-hsa-151136 | 8.48 | 21.54 | 14.45 | 16.73 | 5.39 | 11.68 | 3.21 | 8.13 | 0.70 |
| 46 | piR-hsa-214132 | 12.72 | 23.08 | 19.27 | 8.37 | 12.14 | 13.62 | 0.00 | 0.45 | 0.00 |
| 47 | piR-hsa-1332287 | 6.89 | 52.32 | 13.25 | 0.00 | 5.39 | 5.84 | 0.00 | 2.26 | 1.39 |
| 48 | piR-hsa-1923208 | 6.89 | 7.69 | 2.41 | 3.35 | 2.70 | 1.95 | 35.29 | 5.87 | 19.52 |
| 49 | piR-hsa-1463989 | 2.65 | 21.54 | 27.70 | 6.69 | 5.39 | 19.46 | 0.00 | 0.00 | 0.00 |
| 50 | piR-hsa-2413094 | 12.72 | 30.77 | 20.47 | 0.84 | 10.79 | 7.78 | 0.00 | 0.00 | 0.00 |
| 51 | piR-hsa-2299252 | 14.84 | 4.62 | 28.90 | 10.04 | 8.09 | 15.57 | 0.00 | 0.00 | 0.00 |
| 52 | piR-hsa-2831593 | 6.89 | 1.54 | 27.70 | 11.71 | 14.83 | 1.95 | 0.00 | 11.75 | 5.58 |
| 53 | piR-hsa-1429070 | 2.12 | 4.62 | 7.23 | 3.35 | 6.74 | 56.43 | 0.00 | 0.00 | 0.00 |
| 54 | piR-hsa-2268195 | 14.84 | 4.62 | 32.52 | 1.67 | 4.05 | 13.62 | 1.60 | 1.36 | 4.18 |
| 55 | piR-hsa-4020841 | 5.83 | 23.08 | 30.11 | 1.67 | 5.39 | 11.68 | 0.00 | 0.00 | 0.00 |
| 56 | piR-hsa-1647694 | 9.54 | 12.31 | 30.11 | 8.37 | 6.74 | 7.78 | 0.00 | 0.00 | 1.39 |
| 57 | piR-hsa-359160_2 | 2.12 | 7.69 | 3.61 | 25.10 | 8.09 | 9.73 | 1.60 | 8.59 | 9.06 |
| 58 | piR-hsa-1001021 | 4.77 | 4.62 | 1.20 | 23.42 | 6.74 | 17.51 | 11.23 | 0.00 | 5.58 |
| 59 | piR-hsa-772699 | 0.00 | 0.00 | 0.00 | 53.54 | 8.09 | 7.78 | 3.21 | 0.90 | 1.39 |

| | piRNA | psc1 | psc2 | psc3 | mpc1 | mpc2 | mpc3 | cpc1 | cpc2 | cpc3 |
|---|---|---|---|---|---|---|---|---|---|---|
| 60 | piR-hsa-3916570 | 15.90 | 7.69 | 28.90 | 11.71 | 6.74 | 3.89 | 0.00 | 0.00 | 0.00 |
| 61 | piR-hsa-1916259 | 3.18 | 3.08 | 1.20 | 23.42 | 8.09 | 3.89 | 25.67 | 4.52 | 0.70 |
| 62 | piR-hsa-4322932 | 6.89 | 24.62 | 28.90 | 4.18 | 4.05 | 3.89 | 0.00 | 0.45 | 0.70 |
| 63 | piR-hsa-2521457 | 3.71 | 4.62 | 3.61 | 1.67 | 4.05 | 0.00 | 41.71 | 5.42 | 6.97 |
| 64 | piR-hsa-1903779 | 1.59 | 3.08 | 7.23 | 4.18 | 6.74 | 42.81 | 4.81 | 0.00 | 0.70 |
| 65 | piR-hsa-2646470 | 0.53 | 4.62 | 8.43 | 22.59 | 21.58 | 3.89 | 1.60 | 2.26 | 4.88 |
| 66 | piR-hsa-2497478 | 24.38 | 4.62 | 8.43 | 1.67 | 5.39 | 0.00 | 19.25 | 2.26 | 3.49 |
| 67 | piR-hsa-1875212 | 4.24 | 3.08 | 1.20 | 0.00 | 4.05 | 0.00 | 38.50 | 8.13 | 9.76 |
| 68 | piR-hsa-4387218 | 0.00 | 0.00 | 0.00 | 0.84 | 1.35 | 0.00 | 4.81 | 27.11 | 31.38 |
| 69 | piR-hsa-6913457 | 10.60 | 16.93 | 7.23 | 20.91 | 5.39 | 1.95 | 1.60 | 0.00 | 0.00 |
| 70 | piR-hsa-1756618 | 2.65 | 0.00 | 3.61 | 19.24 | 21.58 | 3.89 | 9.62 | 0.45 | 3.49 |
| 71 | piR-hsa-2592846 | 0.00 | 0.00 | 0.00 | 0.00 | 0.00 | 0.00 | 60.96 | 0.90 | 2.09 |
| 72 | piR-hsa-1908839 | 6.36 | 0.00 | 2.41 | 1.67 | 17.53 | 0.00 | 16.04 | 10.84 | 7.67 |
| 73 | piR-hsa-1576285 | 0.00 | 3.08 | 0.00 | 1.67 | 1.35 | 13.62 | 3.21 | 20.33 | 18.83 |
| 74 | piR-hsa-1862211 | 0.00 | 7.69 | 0.00 | 12.55 | 0.00 | 21.40 | 11.23 | 3.61 | 2.79 |
| 75 | piR-hsa-3658742 | 6.89 | 1.54 | 4.82 | 0.00 | 24.27 | 0.00 | 16.04 | 2.26 | 2.79 |
| 76 | piR-hsa-1872235 | 6.89 | 3.08 | 9.63 | 0.00 | 8.09 | 1.95 | 16.04 | 7.23 | 4.88 |
| 77 | piR-hsa-1277994 | 3.71 | 0.00 | 8.43 | 2.51 | 29.67 | 1.95 | 4.81 | 0.00 | 6.28 |
| 78 | piR-hsa-343382 | 4.24 | 4.62 | 2.41 | 1.67 | 4.05 | 1.95 | 19.25 | 9.04 | 9.76 |
| 79 | piR-hsa-2027369 | 14.84 | 16.93 | 12.04 | 4.18 | 2.70 | 5.84 | 0.00 | 0.00 | 0.00 |
| 80 | piR-hsa-1399886 | 2.12 | 24.62 | 4.82 | 15.89 | 8.09 | 0.00 | 0.00 | 0.00 | 0.00 |
| 81 | piR-hsa-1798104 | 7.42 | 15.39 | 24.09 | 2.51 | 1.35 | 3.89 | 0.00 | 0.00 | 0.00 |
| 82 | piR-hsa-1875155 | 2.12 | 21.54 | 21.68 | 3.35 | 0.00 | 5.84 | 0.00 | 0.00 | 0.00 |
| 83 | piR-hsa-1548068 | 1.59 | 6.15 | 13.25 | 4.18 | 10.79 | 0.00 | 1.60 | 8.13 | 8.37 |
| 84 | piR-hsa-721859_9 | 2.12 | 9.23 | 8.43 | 6.69 | 4.05 | 17.51 | 3.21 | 0.90 | 1.39 |
| 85 | piR-hsa-3732088 | 16.43 | 3.08 | 20.47 | 1.67 | 1.35 | 1.95 | 0.00 | 3.16 | 4.88 |
| 86 | piR-hsa-1303811 | 1.59 | 4.62 | 0.00 | 31.79 | 1.35 | 3.89 | 9.62 | 0.00 | 0.00 |
| 87 | piR-hsa-2481097 | 6.89 | 3.08 | 6.02 | 0.00 | 12.14 | 0.00 | 19.25 | 4.07 | 1.39 |
| 88 | piR-hsa-4028152 | 2.12 | 15.39 | 0.00 | 5.02 | 2.70 | 25.30 | 1.60 | 0.00 | 0.70 |
| 89 | piR-hsa-389007 | 2.65 | 4.62 | 2.41 | 9.20 | 10.79 | 19.46 | 3.21 | 0.00 | 0.00 |
| 90 | piR-hsa-2477264 | 5.30 | 6.15 | 6.02 | 0.00 | 10.79 | 0.00 | 17.64 | 4.07 | 2.09 |
| 91 | piR-hsa-255984 | 2.12 | 3.08 | 12.04 | 5.86 | 2.70 | 23.35 | 0.00 | 0.00 | 2.09 |
| 92 | piR-hsa-137136 | 7.42 | 7.69 | 9.63 | 8.37 | 8.09 | 9.73 | 0.00 | 0.00 | 0.00 |
| 93 | piR-hsa-3807498 | 7.95 | 13.85 | 15.66 | 1.67 | 6.74 | 3.89 | 0.00 | 0.00 | 0.70 |
| 94 | piR-hsa-783698 | 15.37 | 7.69 | 6.02 | 5.02 | 13.49 | 1.95 | 0.00 | 0.00 | 0.00 |
| 95 | piR-hsa-1988800 | 7.95 | 4.62 | 12.04 | 3.35 | 14.83 | 5.84 | 0.00 | 0.00 | 0.00 |
| 96 | piR-hsa-1941780 | 3.71 | 6.15 | 3.61 | 1.67 | 2.70 | 1.95 | 19.25 | 1.36 | 7.67 |
| 97 | piR-hsa-3875265 | 10.60 | 9.23 | 10.84 | 5.86 | 9.44 | 1.95 | 0.00 | 0.00 | 0.00 |
| 98 | piR-hsa-1876265 | 4.24 | 4.62 | 3.61 | 0.00 | 0.00 | 0.00 | 19.25 | 6.33 | 9.76 |
| 99 | piR-hsa-368987 | 6.36 | 4.62 | 4.82 | 5.02 | 4.05 | 3.89 | 9.62 | 5.87 | 3.49 |
| 100 | piR-hsa-1890632 | 6.36 | 4.62 | 7.23 | 0.00 | 5.39 | 0.00 | 11.23 | 4.52 | 8.37 |
| 101 | piR-hsa-4078407 | 2.12 | 20.00 | 9.63 | 10.04 | 1.35 | 3.89 | 0.00 | 0.00 | 0.00 |
| 102 | piR-hsa-76848 | 4.24 | 4.62 | 1.20 | 1.67 | 1.35 | 25.30 | 1.60 | 1.36 | 5.58 |
| 103 | piR-hsa-2565910 | 0.53 | 12.31 | 4.82 | 5.02 | 5.39 | 15.57 | 0.00 | 1.81 | 1.39 |
| 104 | piR-hsa-1205256 | 3.18 | 4.62 | 1.20 | 21.75 | 0.00 | 11.68 | 3.21 | 0.45 | 0.00 |
| 105 | piR-hsa-2840936 | 10.60 | 0.00 | 1.20 | 0.00 | 6.74 | 0.00 | 19.25 | 4.97 | 2.79 |
| 106 | piR-hsa-20628 | 3.71 | 9.23 | 12.04 | 2.51 | 8.09 | 9.73 | 0.00 | 0.00 | 0.00 |
| 107 | piR-hsa-753191 | 0.00 | 0.00 | 0.00 | 0.00 | 0.00 | 0.00 | 35.29 | 5.42 | 4.18 |
| 108 | piR-hsa-3683883 | 2.12 | 3.08 | 6.02 | 20.08 | 1.35 | 1.95 | 6.42 | 0.00 | 2.79 |
| 109 | piR-hsa-1690788 | 1.59 | 4.62 | 4.82 | 5.86 | 6.74 | 17.51 | 0.00 | 0.00 | 2.09 |
| 110 | piR-hsa-2530015 | 6.89 | 3.08 | 1.20 | 2.51 | 0.00 | 0.00 | 16.04 | 2.26 | 11.16 |
| 111 | piR-hsa-346271 | 3.71 | 3.08 | 15.66 | 1.67 | 2.70 | 0.00 | 3.21 | 1.81 | 10.46 |
| 112 | piR-hsa-3546008 | 6.89 | 3.08 | 8.43 | 5.02 | 14.83 | 3.89 | 0.00 | 0.00 | 0.00 |
| 113 | piR-hsa-5982715 | 7.42 | 13.85 | 2.41 | 6.69 | 2.70 | 7.78 | 0.00 | 0.90 | 0.00 |
| 114 | piR-hsa-2398570 | 1.06 | 6.15 | 3.61 | 2.51 | 12.14 | 15.57 | 0.00 | 0.00 | 0.70 |
| 115 | piR-hsa-1883972 | 6.36 | 0.00 | 0.00 | 0.00 | 2.70 | 0.00 | 19.25 | 3.61 | 9.76 |
| 116 | piR-hsa-3718263_3 | 0.00 | 0.00 | 0.00 | 0.00 | 0.00 | 0.00 | 33.69 | 5.87 | 2.09 |
| 117 | piR-hsa-4472891 | 2.12 | 3.08 | 9.63 | 11.71 | 6.74 | 5.84 | 0.00 | 1.81 | 0.70 |
| 118 | piR-hsa-2208850 | 4.24 | 15.39 | 7.23 | 5.02 | 1.35 | 7.78 | 0.00 | 0.00 | 0.00 |

| | piRNA | psc1 | psc2 | psc3 | mpc1 | mpc2 | mpc3 | cpc1 | cpc2 | cpc3 |
|---|---|---|---|---|---|---|---|---|---|---|
| 119 | piR-hsa-785969 | 2.12 | 3.08 | 0.00 | 6.69 | 0.00 | 1.95 | 24.06 | 1.36 | 1.39 |
| 120 | piR-hsa-163695 | 6.36 | 10.77 | 7.23 | 0.84 | 6.74 | 7.78 | 0.00 | 0.00 | 0.70 |
| 121 | piR-hsa-2490897 | 6.89 | 0.00 | 1.20 | 1.67 | 0.00 | 0.00 | 11.23 | 5.42 | 13.95 |
| 122 | piR-hsa-2151268 | 1.59 | 7.69 | 3.61 | 7.53 | 8.09 | 11.68 | 0.00 | 0.00 | 0.00 |
| 123 | piR-hsa-771714_3 | 3.71 | 4.62 | 2.41 | 0.00 | 0.00 | 3.89 | 17.64 | 0.90 | 6.97 |
| 124 | piR-hsa-1920687 | 4.24 | 4.62 | 1.20 | 1.67 | 4.05 | 0.00 | 12.83 | 5.42 | 5.58 |
| 125 | piR-hsa-2351941 | 5.30 | 23.08 | 2.41 | 2.51 | 2.70 | 1.95 | 1.60 | 0.00 | 0.00 |
| 126 | piR-hsa-3265318 | 3.18 | 13.85 | 6.02 | 1.67 | 2.70 | 11.68 | 0.00 | 0.00 | 0.00 |
| 127 | piR-hsa-2426792 | 2.12 | 24.62 | 4.82 | 6.69 | 0.00 | 0.00 | 0.00 | 0.00 | 0.70 |
| 128 | piR-hsa-2503702 | 13.25 | 0.00 | 8.43 | 0.00 | 4.05 | 0.00 | 6.42 | 3.16 | 2.79 |
| 129 | piR-hsa-161264_3 | 1.06 | 0.00 | 1.20 | 8.37 | 5.39 | 21.40 | 0.00 | 0.00 | 0.00 |
| 130 | piR-hsa-1912443 | 1.59 | 0.00 | 1.20 | 0.00 | 1.35 | 0.00 | 28.87 | 2.26 | 2.09 |
| 131 | piR-hsa-2230204 | 8.48 | 6.15 | 15.66 | 3.35 | 2.70 | 0.00 | 0.00 | 0.00 | 0.70 |
| 132 | piR-hsa-2090264 | 7.42 | 1.54 | 25.29 | 0.84 | 1.35 | 0.00 | 0.00 | 0.00 | 0.00 |
| 133 | piR-hsa-1242358 | 0.00 | 0.00 | 0.00 | 0.00 | 0.00 | 0.00 | 32.08 | 2.26 | 2.09 |
| 134 | piR-hsa-724912 | 2.65 | 1.54 | 3.61 | 22.59 | 1.35 | 0.00 | 3.21 | 0.45 | 0.70 |
| 135 | piR-hsa-7765828 | 1.06 | 7.69 | 3.61 | 13.38 | 4.05 | 5.84 | 0.00 | 0.00 | 0.00 |
| 136 | piR-hsa-2172087 | 1.59 | 10.77 | 3.61 | 5.02 | 2.70 | 7.78 | 3.21 | 0.90 | 0.00 |
| 137 | piR-hsa-1425899 | 0.00 | 4.62 | 1.20 | 6.69 | 2.70 | 17.51 | 0.00 | 1.36 | 1.39 |
| 138 | piR-hsa-1921188 | 1.59 | 3.08 | 1.20 | 0.00 | 10.79 | 0.00 | 14.44 | 3.61 | 0.70 |
| 139 | piR-hsa-1917139 | 2.65 | 1.54 | 1.20 | 0.00 | 4.05 | 0.00 | 20.85 | 1.36 | 3.49 |
| 140 | piR-hsa-3829948 | 0.53 | 3.08 | 0.00 | 13.38 | 4.05 | 13.62 | 0.00 | 0.00 | 0.00 |
| 141 | piR-hsa-3735787 | 7.42 | 3.08 | 2.41 | 0.00 | 2.70 | 0.00 | 12.83 | 4.07 | 2.09 |
| 142 | piR-hsa-5996985 | 0.00 | 0.00 | 0.00 | 0.00 | 0.00 | 0.00 | 28.87 | 5.42 | 0.00 |
| 143 | piR-hsa-2989729 | 6.36 | 6.15 | 13.25 | 0.84 | 2.70 | 3.89 | 0.00 | 0.00 | 0.00 |
| 144 | piR-hsa-21839 | 0.53 | 3.08 | 0.00 | 24.26 | 2.70 | 1.95 | 0.00 | 0.45 | 0.00 |
| 145 | piR-hsa-1872463 | 0.00 | 4.62 | 1.20 | 1.67 | 0.00 | 19.46 | 0.00 | 3.16 | 2.79 |
| 146 | piR-hsa-508592 | 2.12 | 13.85 | 7.23 | 2.51 | 1.35 | 5.84 | 0.00 | 0.00 | 0.00 |
| 147 | piR-hsa-2252211 | 1.59 | 6.15 | 3.61 | 0.84 | 20.23 | 0.00 | 0.00 | 0.45 | 0.00 |
| 148 | piR-hsa-4110708 | 2.65 | 27.70 | 1.20 | 0.84 | 0.00 | 0.00 | 0.00 | 0.00 | 0.00 |
| 149 | piR-hsa-298158 | 4.24 | 6.15 | 6.02 | 7.53 | 4.05 | 3.89 | 0.00 | 0.00 | 0.00 |
| 150 | piR-hsa-7892960 | 6.36 | 1.54 | 3.61 | 4.18 | 2.70 | 1.95 | 3.21 | 4.52 | 3.49 |
| 151 | piR-hsa-2395910 | 2.65 | 3.08 | 3.61 | 3.35 | 1.35 | 17.51 | 0.00 | 0.00 | 0.00 |
| 152 | piR-hsa-2536290 | 0.53 | 12.31 | 3.61 | 8.37 | 1.35 | 3.89 | 0.00 | 0.45 | 0.70 |
| 153 | piR-hsa-2152778 | 2.65 | 10.77 | 6.02 | 5.02 | 2.70 | 3.89 | 0.00 | 0.00 | 0.00 |
| 154 | piR-hsa-5077723 | 0.53 | 4.62 | 12.04 | 8.37 | 5.39 | 0.00 | 0.00 | 0.00 | 0.00 |
| 155 | piR-hsa-1291516 | 7.42 | 6.15 | 10.84 | 2.51 | 1.35 | 0.00 | 0.00 | 0.90 | 1.39 |
| 156 | piR-hsa-1901970 | 12.19 | 7.69 | 0.00 | 3.35 | 5.39 | 1.95 | 0.00 | 0.00 | 0.00 |
| 157 | piR-hsa-2450089_2 | 0.00 | 1.54 | 4.82 | 3.35 | 1.35 | 19.46 | 0.00 | 0.00 | 0.00 |
| 158 | piR-hsa-108574 | 3.71 | 7.69 | 4.82 | 3.35 | 8.09 | 1.95 | 0.00 | 0.00 | 0.70 |
| 159 | piR-hsa-1259653 | 2.65 | 1.54 | 2.41 | 0.84 | 1.35 | 1.95 | 16.04 | 1.81 | 0.70 |
| 160 | piR-hsa-3136454 | 2.65 | 16.93 | 2.41 | 1.67 | 2.70 | 1.95 | 0.00 | 0.00 | 0.70 |
| 161 | piR-hsa-2286229 | 4.24 | 3.08 | 9.63 | 4.18 | 5.39 | 1.95 | 0.00 | 0.45 | 0.00 |
| 162 | piR-hsa-2209630 | 2.65 | 0.00 | 0.00 | 23.42 | 2.70 | 0.00 | 0.00 | 0.00 | 0.00 |
| 163 | piR-hsa-1696540 | 0.00 | 0.00 | 0.00 | 0.00 | 0.00 | 1.95 | 24.06 | 1.81 | 0.70 |
| 164 | piR-hsa-6245615 | 3.71 | 4.62 | 7.23 | 0.00 | 10.79 | 1.95 | 0.00 | 0.00 | 0.00 |
| 165 | piR-hsa-2072163 | 1.06 | 4.62 | 2.41 | 1.67 | 2.70 | 5.84 | 8.02 | 1.81 | 0.00 |
| 166 | piR-hsa-1557538 | 0.00 | 0.00 | 2.41 | 8.37 | 9.44 | 7.78 | 0.00 | 0.00 | 0.00 |
| 167 | piR-hsa-2844156 | 7.42 | 3.08 | 4.82 | 5.86 | 0.00 | 3.89 | 0.00 | 1.36 | 1.39 |
| 168 | piR-hsa-6744266 | 0.00 | 1.54 | 1.20 | 5.86 | 0.00 | 5.84 | 9.62 | 2.26 | 1.39 |
| 169 | piR-hsa-1882039 | 1.06 | 0.00 | 0.00 | 0.00 | 0.00 | 0.00 | 17.64 | 2.71 | 6.28 |
| 170 | piR-hsa-2137611 | 5.30 | 15.39 | 6.02 | 0.84 | 0.00 | 0.00 | 0.00 | 0.00 | 0.00 |
| 171 | piR-hsa-58291 | 3.71 | 6.15 | 8.43 | 0.84 | 4.05 | 3.89 | 0.00 | 0.45 | 0.00 |
| 172 | piR-hsa-4408495 | 0.00 | 0.00 | 2.41 | 5.86 | 5.39 | 11.68 | 1.60 | 0.45 | 0.00 |
| 173 | piR-hsa-374600 | 1.06 | 0.00 | 1.20 | 10.87 | 1.35 | 0.00 | 12.83 | 0.00 | 0.00 |
| 174 | piR-hsa-2829712 | 3.71 | 0.00 | 1.20 | 0.00 | 5.39 | 0.00 | 9.62 | 3.16 | 4.18 |
| 175 | piR-hsa-3177742 | 3.18 | 4.62 | 7.23 | 1.67 | 0.00 | 9.73 | 0.00 | 0.00 | 0.70 |
| 176 | piR-hsa-1585146 | 2.12 | 0.00 | 4.82 | 9.20 | 6.74 | 3.89 | 0.00 | 0.00 | 0.00 |
| 177 | piR-hsa-4403577 | 2.65 | 4.62 | 7.23 | 0.84 | 5.39 | 3.89 | 0.00 | 1.36 | 0.70 |

| | piRNA | psc1 | psc2 | psc3 | mpc1 | mpc2 | mpc3 | cpc1 | cpc2 | cpc3 |
|---|---|---|---|---|---|---|---|---|---|---|
| 178 | piR-hsa-1528884 | 4.24 | 10.77 | 4.82 | 2.51 | 1.35 | 1.95 | 0.00 | 0.00 | 0.70 |
| 179 | piR-hsa-2494226 | 4.77 | 3.08 | 0.00 | 0.00 | 2.70 | 0.00 | 8.02 | 2.71 | 4.88 |
| 180 | piR-hsa-229786 | 0.00 | 4.62 | 0.00 | 2.51 | 2.70 | 15.57 | 0.00 | 0.00 | 0.70 |
| 181 | piR-hsa-1492262 | 3.71 | 4.62 | 1.20 | 0.84 | 12.14 | 0.00 | 1.60 | 0.45 | 1.39 |
| 182 | piR-hsa-132896 | 1.59 | 4.62 | 4.82 | 4.18 | 8.09 | 1.95 | 0.00 | 0.00 | 0.70 |
| 183 | piR-hsa-1376916 | 2.12 | 10.77 | 2.41 | 3.35 | 2.70 | 3.89 | 0.00 | 0.45 | 0.00 |
| 184 | piR-hsa-160969 | 0.53 | 0.00 | 6.02 | 3.35 | 2.70 | 11.68 | 0.00 | 0.00 | 1.39 |
| 185 | piR-hsa-768321 | 8.48 | 3.08 | 1.20 | 0.84 | 0.00 | 0.00 | 4.81 | 0.90 | 6.28 |
| 186 | piR-hsa-3842249 | 1.59 | 0.00 | 2.41 | 10.04 | 1.35 | 5.84 | 3.21 | 0.45 | 0.70 |
| 187 | piR-hsa-362913 | 1.06 | 1.54 | 1.20 | 21.75 | 0.00 | 0.00 | 0.00 | 0.00 | 0.00 |
| 188 | piR-hsa-1870459 | 3.18 | 0.00 | 4.82 | 0.84 | 6.74 | 0.00 | 4.81 | 2.71 | 2.09 |
| 189 | piR-hsa-3634880 | 2.12 | 0.00 | 0.00 | 5.02 | 6.74 | 3.89 | 3.21 | 0.00 | 4.18 |
| 190 | piR-hsa-1941637 | 2.65 | 0.00 | 0.00 | 16.73 | 1.35 | 0.00 | 3.21 | 0.45 | 0.70 |
| 191 | piR-hsa-834074 | 0.00 | 0.00 | 0.00 | 0.00 | 0.00 | 0.00 | 20.85 | 2.71 | 1.39 |
| 192 | piR-hsa-1900529 | 5.83 | 3.08 | 3.61 | 0.00 | 4.05 | 1.95 | 0.00 | 2.26 | 4.18 |
| 193 | piR-hsa-2220917 | 0.00 | 3.08 | 1.20 | 11.71 | 8.09 | 0.00 | 0.00 | 0.00 | 0.70 |
| 194 | piR-hsa-3739406 | 3.18 | 1.54 | 2.41 | 5.86 | 2.70 | 0.00 | 1.60 | 0.45 | 6.97 |
| 195 | piR-hsa-7308134 | 0.00 | 0.00 | 0.00 | 14.22 | 2.70 | 7.78 | 0.00 | 0.00 | 0.00 |
| 196 | piR-hsa-1708978 | 0.53 | 0.00 | 0.00 | 11.71 | 2.70 | 9.73 | 0.00 | 0.00 | 0.00 |
| 197 | piR-hsa-3231825 | 3.71 | 12.31 | 3.61 | 1.67 | 1.35 | 1.95 | 0.00 | 0.00 | 0.00 |
| 198 | piR-hsa-2398119 | 4.77 | 3.08 | 0.00 | 1.67 | 5.39 | 0.00 | 9.62 | 0.00 | 0.00 |
| 199 | piR-hsa-2882083 | 3.71 | 4.62 | 0.00 | 5.86 | 4.05 | 1.95 | 1.60 | 1.36 | 1.39 |
| 200 | piR-hsa-315964 | 4.77 | 1.54 | 10.84 | 1.67 | 5.39 | 0.00 | 0.00 | 0.00 | 0.00 |
| 201 | piR-hsa-4030155 | 6.89 | 4.62 | 2.41 | 1.67 | 2.70 | 5.84 | 0.00 | 0.00 | 0.00 |
| 202 | piR-hsa-2615134 | 3.71 | 3.08 | 2.41 | 10.87 | 4.05 | 0.00 | 0.00 | 0.00 | 0.00 |
| 203 | piR-hsa-7544198 | 1.59 | 0.00 | 2.41 | 12.55 | 6.74 | 0.00 | 0.00 | 0.00 | 0.70 |
| 204 | piR-hsa-1843231 | 1.06 | 4.62 | 4.82 | 4.18 | 5.39 | 3.89 | 0.00 | 0.00 | 0.00 |
| 205 | piR-hsa-2670375 | 1.06 | 1.54 | 2.41 | 11.71 | 1.35 | 5.84 | 0.00 | 0.00 | 0.00 |
| 206 | piR-hsa-2464166 | 1.06 | 3.08 | 0.00 | 0.84 | 1.35 | 17.51 | 0.00 | 0.00 | 0.00 |
| 207 | piR-hsa-2589139 | 1.59 | 1.54 | 4.82 | 12.55 | 1.35 | 1.95 | 0.00 | 0.00 | 0.00 |
| 208 | piR-hsa-3119265 | 4.24 | 1.54 | 4.82 | 2.51 | 6.74 | 3.89 | 0.00 | 0.00 | 0.00 |
| 209 | piR-hsa-1531418 | 1.06 | 3.08 | 3.61 | 6.69 | 5.39 | 3.89 | 0.00 | 0.00 | 0.00 |
| 210 | piR-hsa-2253283 | 2.65 | 4.62 | 0.00 | 11.71 | 2.70 | 1.95 | 0.00 | 0.00 | 0.00 |
| 211 | piR-hsa-2148238 | 1.59 | 7.69 | 8.43 | 2.51 | 1.35 | 1.95 | 0.00 | 0.00 | 0.00 |
| 212 | piR-hsa-1595580 | 2.12 | 6.15 | 7.23 | 2.51 | 5.39 | 0.00 | 0.00 | 0.00 | 0.00 |
| 213 | piR-hsa-3978322 | 0.00 | 0.00 | 0.00 | 10.04 | 6.74 | 3.89 | 1.60 | 0.90 | 0.00 |
| 214 | piR-hsa-1773241 | 2.65 | 1.54 | 3.61 | 3.35 | 8.09 | 3.89 | 0.00 | 0.00 | 0.00 |
| 215 | piR-hsa-1434629 | 0.53 | 1.54 | 3.61 | 7.53 | 4.05 | 5.84 | 0.00 | 0.00 | 0.00 |
| 216 | piR-hsa-1340768 | 0.53 | 1.54 | 1.20 | 3.35 | 6.74 | 7.78 | 0.00 | 0.45 | 1.39 |
| 217 | piR-hsa-333507 | 1.59 | 4.62 | 8.43 | 1.67 | 2.70 | 3.89 | 0.00 | 0.00 | 0.00 |
| 218 | piR-hsa-118348 | 3.18 | 6.15 | 3.61 | 3.35 | 4.05 | 0.00 | 0.00 | 0.45 | 2.09 |
| 219 | piR-hsa-3623001 | 4.24 | 1.54 | 1.20 | 0.00 | 9.44 | 0.00 | 1.60 | 1.36 | 3.49 |
| 220 | piR-hsa-2090890 | 2.65 | 4.62 | 4.82 | 4.18 | 2.70 | 3.89 | 0.00 | 0.00 | 0.00 |
| 221 | piR-hsa-1929067 | 2.12 | 9.23 | 1.20 | 2.51 | 0.00 | 7.78 | 0.00 | 0.00 | 0.00 |
| 222 | piR-hsa-2827579 | 1.06 | 0.00 | 0.00 | 0.00 | 1.35 | 1.95 | 6.42 | 0.90 | 11.16 |
| 223 | piR-hsa-1707103 | 0.00 | 1.54 | 0.00 | 0.00 | 0.00 | 0.00 | 16.04 | 4.52 | 0.70 |
| 224 | piR-hsa-1256360 | 0.00 | 0.00 | 0.00 | 9.20 | 2.70 | 9.73 | 0.00 | 0.45 | 0.70 |
| 225 | piR-hsa-1919455 | 6.89 | 1.54 | 2.41 | 0.00 | 1.35 | 0.00 | 4.81 | 3.61 | 2.09 |
| 226 | piR-hsa-1607096 | 1.06 | 12.31 | 1.20 | 0.84 | 2.70 | 3.89 | 0.00 | 0.00 | 0.70 |
| 227 | piR-hsa-2515454 | 5.30 | 3.08 | 0.00 | 0.00 | 5.39 | 0.00 | 6.42 | 1.81 | 0.70 |
| 228 | piR-hsa-368381 | 0.53 | 1.54 | 1.20 | 2.51 | 0.00 | 1.95 | 4.81 | 3.16 | 6.97 |
| 229 | piR-hsa-4379982 | 4.24 | 4.62 | 1.20 | 0.84 | 9.44 | 0.00 | 1.60 | 0.00 | 0.70 |
| 230 | piR-hsa-2240007 | 3.71 | 7.69 | 4.82 | 1.67 | 2.70 | 1.95 | 0.00 | 0.00 | 0.00 |
| 231 | piR-hsa-4416099_9 | 0.00 | 0.00 | 1.20 | 0.84 | 0.00 | 19.46 | 0.00 | 0.90 | 0.00 |
| 232 | piR-hsa-7106256 | 1.59 | 0.00 | 1.20 | 10.87 | 5.39 | 1.95 | 0.00 | 0.45 | 0.70 |
| 233 | piR-hsa-1481120 | 2.12 | 3.08 | 1.20 | 8.37 | 5.39 | 1.95 | 0.00 | 0.00 | 0.00 |
| 234 | piR-hsa-1284504 | 0.53 | 1.54 | 0.00 | 4.18 | 0.00 | 3.89 | 6.42 | 2.71 | 2.79 |
| 235 | piR-hsa-2505515 | 0.00 | 0.00 | 0.00 | 7.53 | 6.74 | 7.78 | 0.00 | 0.00 | 0.00 |
| 236 | piR-hsa-1927627 | 0.00 | 0.00 | 0.00 | 0.00 | 0.00 | 0.00 | 16.04 | 1.81 | 4.18 |

| | piRNA | psc1 | psc2 | psc3 | mpc1 | mpc2 | mpc3 | cpc1 | cpc2 | cpc3 |
|---|---|---|---|---|---|---|---|---|---|---|
| 237 | piR-hsa-1921551 | 2.12 | 1.54 | 1.20 | 12.55 | 2.70 | 0.00 | 0.00 | 0.45 | 1.39 |
| 238 | piR-hsa-163499 | 4.77 | 3.08 | 3.61 | 1.67 | 2.70 | 1.95 | 1.60 | 0.45 | 2.09 |
| 239 | piR-hsa-4202081 | 1.59 | 1.54 | 4.82 | 4.18 | 5.39 | 3.89 | 0.00 | 0.45 | 0.00 |
| 240 | piR-hsa-147696 | 1.59 | 0.00 | 0.00 | 0.84 | 1.35 | 0.00 | 9.62 | 4.07 | 4.18 |
| 241 | piR-hsa-2353109 | 0.00 | 12.31 | 1.20 | 0.84 | 1.35 | 5.84 | 0.00 | 0.00 | 0.00 |
| 242 | piR-hsa-8270846 | 3.71 | 4.62 | 7.23 | 5.86 | 0.00 | 0.00 | 0.00 | 0.00 | 0.00 |
| 243 | piR-hsa-1905680 | 0.53 | 1.54 | 3.61 | 0.84 | 0.00 | 0.00 | 8.02 | 4.07 | 2.79 |
| 244 | piR-hsa-307961 | 2.12 | 6.15 | 0.00 | 3.35 | 0.00 | 9.73 | 0.00 | 0.00 | 0.00 |
| 245 | piR-hsa-2333057 | 4.77 | 0.00 | 3.61 | 0.84 | 9.44 | 1.95 | 0.00 | 0.00 | 0.70 |
| 246 | piR-hsa-1938524 | 0.53 | 0.00 | 1.20 | 0.84 | 1.35 | 1.95 | 11.23 | 1.36 | 2.79 |
| 247 | piR-hsa-3232943 | 3.18 | 0.00 | 3.61 | 5.02 | 5.39 | 3.89 | 0.00 | 0.00 | 0.00 |
| 248 | piR-hsa-3513154 | 5.30 | 1.54 | 7.23 | 4.18 | 2.70 | 0.00 | 0.00 | 0.00 | 0.00 |
| 249 | piR-hsa-3674332 | 14.31 | 0.00 | 0.00 | 0.00 | 2.70 | 0.00 | 1.60 | 0.90 | 1.39 |
| 250 | piR-hsa-2213434 | 6.36 | 3.08 | 7.23 | 4.18 | 0.00 | 0.00 | 0.00 | 0.00 | 0.00 |
| 251 | piR-hsa-2308163 | 2.65 | 7.69 | 3.61 | 4.18 | 2.70 | 0.00 | 0.00 | 0.00 | 0.00 |
| 252 | piR-hsa-1748898 | 0.00 | 7.69 | 4.82 | 5.02 | 1.35 | 1.95 | 0.00 | 0.00 | 0.00 |
| 253 | piR-hsa-2525461 | 0.00 | 3.08 | 3.61 | 6.69 | 2.70 | 1.95 | 0.00 | 0.00 | 2.79 |
| 254 | piR-hsa-1296118 | 3.18 | 0.00 | 2.41 | 0.00 | 2.70 | 0.00 | 8.02 | 3.61 | 0.70 |
| 255 | piR-hsa-1632961 | 0.53 | 12.31 | 3.61 | 0.84 | 1.35 | 1.95 | 0.00 | 0.00 | 0.00 |
| 256 | piR-hsa-728085 | 0.53 | 1.54 | 0.00 | 0.84 | 0.00 | 0.00 | 17.64 | 0.00 | 0.00 |
| 257 | piR-hsa-623353 | 0.00 | 1.54 | 0.00 | 0.84 | 0.00 | 15.57 | 0.00 | 0.45 | 2.09 |
| 258 | piR-hsa-3527815 | 1.06 | 13.85 | 3.61 | 0.00 | 0.00 | 1.95 | 0.00 | 0.00 | 0.00 |
| 259 | piR-hsa-2478880_2 | 0.00 | 1.54 | 0.00 | 0.00 | 2.70 | 0.00 | 8.02 | 3.16 | 4.88 |
| 260 | piR-hsa-3558751 | 4.24 | 6.15 | 4.82 | 1.67 | 1.35 | 1.95 | 0.00 | 0.00 | 0.00 |
| 261 | piR-hsa-4178299 | 5.30 | 4.62 | 7.23 | 1.67 | 1.35 | 0.00 | 0.00 | 0.00 | 0.00 |
| 262 | piR-hsa-5411637 | 2.65 | 6.15 | 2.41 | 0.84 | 4.05 | 3.89 | 0.00 | 0.00 | 0.00 |
| 263 | piR-hsa-1883893 | 0.00 | 7.69 | 0.00 | 5.86 | 0.00 | 3.89 | 0.00 | 1.81 | 0.70 |
| 264 | piR-hsa-2742244 | 1.59 | 0.00 | 0.00 | 0.00 | 4.05 | 0.00 | 9.62 | 1.81 | 2.79 |
| 265 | piR-hsa-669874 | 0.53 | 0.00 | 0.00 | 19.24 | 0.00 | 0.00 | 0.00 | 0.00 | 0.00 |
| 266 | piR-hsa-645846 | 3.18 | 4.62 | 3.61 | 0.84 | 1.35 | 5.84 | 0.00 | 0.00 | 0.00 |
| 267 | piR-hsa-3776081 | 0.53 | 1.54 | 0.00 | 9.20 | 8.09 | 0.00 | 0.00 | 0.00 | 0.00 |
| 268 | piR-hsa-211123 | 2.12 | 0.00 | 1.20 | 5.02 | 5.39 | 3.89 | 0.00 | 0.90 | 0.70 |
| 269 | piR-hsa-2248086 | 0.53 | 4.62 | 6.02 | 3.35 | 2.70 | 1.95 | 0.00 | 0.00 | 0.00 |
| 270 | piR-hsa-1909905 | 0.53 | 0.00 | 0.00 | 1.67 | 1.35 | 15.57 | 0.00 | 0.00 | 0.00 |
| 271 | piR-hsa-1593307 | 3.18 | 4.62 | 6.02 | 1.67 | 1.35 | 1.95 | 0.00 | 0.00 | 0.00 |
| 272 | piR-hsa-2042088 | 0.00 | 0.00 | 0.00 | 0.00 | 0.00 | 0.00 | 12.83 | 4.52 | 1.39 |
| 273 | piR-hsa-2490509 | 0.00 | 1.54 | 0.00 | 0.00 | 1.35 | 0.00 | 12.83 | 0.90 | 2.09 |
| 274 | piR-hsa-144277_2 | 0.00 | 0.00 | 0.00 | 0.00 | 5.39 | 0.00 | 11.23 | 1.36 | 0.70 |
| 275 | piR-hsa-4131663 | 0.00 | 0.00 | 0.00 | 4.18 | 4.05 | 1.95 | 8.02 | 0.45 | 0.00 |
| 276 | piR-hsa-2542835 | 1.59 | 0.00 | 1.20 | 0.00 | 2.70 | 0.00 | 11.23 | 0.45 | 1.39 |
| 277 | piR-hsa-1229611 | 2.12 | 1.54 | 1.20 | 4.18 | 2.70 | 5.84 | 0.00 | 0.90 | 0.00 |
| 278 | piR-hsa-2423519 | 1.06 | 0.00 | 0.00 | 0.00 | 4.05 | 11.68 | 1.60 | 0.00 | 0.00 |
| 279 | piR-hsa-3021684 | 4.24 | 1.54 | 3.61 | 7.53 | 1.35 | 0.00 | 0.00 | 0.00 | 0.00 |
| 280 | piR-hsa-2490287 | 0.00 | 1.54 | 0.00 | 3.35 | 1.35 | 1.95 | 6.42 | 2.26 | 1.39 |
| 281 | piR-hsa-1259933 | 1.06 | 1.54 | 0.00 | 8.37 | 2.70 | 0.00 | 1.60 | 2.26 | 0.70 |
| 282 | piR-hsa-1302552 | 0.53 | 0.00 | 0.00 | 5.86 | 6.74 | 0.00 | 0.00 | 0.00 | 4.88 |
| 283 | piR-hsa-67957 | 1.06 | 13.85 | 0.00 | 0.00 | 0.00 | 0.00 | 0.00 | 0.90 | 2.09 |
| 284 | piR-hsa-1726249 | 6.89 | 1.54 | 7.23 | 0.84 | 1.35 | 0.00 | 0.00 | 0.00 | 0.00 |
| 285 | piR-hsa-1409954 | 1.59 | 1.54 | 8.43 | 0.84 | 5.39 | 0.00 | 0.00 | 0.00 | 0.00 |
| 286 | piR-hsa-4053516 | 2.65 | 1.54 | 7.23 | 1.67 | 2.70 | 1.95 | 0.00 | 0.00 | 0.00 |
| 287 | piR-hsa-2425220 | 0.00 | 1.54 | 1.20 | 5.02 | 0.00 | 0.00 | 3.21 | 1.81 | 4.88 |
| 288 | piR-hsa-1686806 | 2.12 | 3.08 | 3.61 | 4.18 | 2.70 | 1.95 | 0.00 | 0.00 | 0.00 |
| 289 | piR-hsa-2319750 | 2.65 | 0.00 | 7.23 | 1.67 | 4.05 | 1.95 | 0.00 | 0.00 | 0.00 |
| 290 | piR-hsa-4397384 | 0.00 | 0.00 | 3.61 | 4.18 | 0.00 | 9.73 | 0.00 | 0.00 | 0.00 |
| 291 | piR-hsa-2829413 | 1.06 | 0.00 | 1.20 | 0.00 | 1.35 | 0.00 | 8.02 | 0.90 | 4.88 |
| 292 | piR-hsa-3839126 | 0.00 | 0.00 | 0.00 | 7.53 | 4.05 | 5.84 | 0.00 | 0.00 | 0.00 |
| 293 | piR-hsa-3944431 | 3.71 | 3.08 | 3.61 | 1.67 | 1.35 | 3.89 | 0.00 | 0.00 | 0.00 |
| 294 | piR-hsa-363100_2 | 2.65 | 1.54 | 0.00 | 1.67 | 4.05 | 0.00 | 3.21 | 1.36 | 2.79 |
| 295 | piR-hsa-6482184 | 0.53 | 10.77 | 1.20 | 0.84 | 0.00 | 3.89 | 0.00 | 0.00 | 0.00 |

| | piRNA | psc1 | psc2 | psc3 | mpc1 | mpc2 | mpc3 | cpc1 | cpc2 | cpc3 |
|---|---|---|---|---|---|---|---|---|---|---|
| 296 | piR-hsa-642866 | 0.00 | 0.00 | 0.00 | 0.00 | 0.00 | 0.00 | 12.83 | 2.26 | 2.09 |
| 297 | piR-hsa-1922210 | 0.53 | 0.00 | 1.20 | 9.20 | 0.00 | 3.89 | 0.00 | 0.90 | 1.39 |
| 298 | piR-hsa-2436454 | 3.18 | 3.08 | 4.82 | 1.67 | 0.00 | 3.89 | 0.00 | 0.45 | 0.00 |
| 299 | piR-hsa-7821967_3 | 0.00 | 0.00 | 1.20 | 0.00 | 0.00 | 0.00 | 11.23 | 3.16 | 1.39 |
| 300 | piR-hsa-4424378 | 0.00 | 0.00 | 0.00 | 0.00 | 1.35 | 0.00 | 12.83 | 1.36 | 1.39 |
| 301 | piR-hsa-237221 | 1.59 | 1.54 | 0.00 | 10.87 | 2.70 | 0.00 | 0.00 | 0.00 | 0.00 |
| 302 | piR-hsa-7833890 | 1.59 | 1.54 | 0.00 | 5.02 | 2.70 | 5.84 | 0.00 | 0.00 | 0.00 |
| 303 | piR-hsa-1905329 | 0.00 | 3.08 | 0.00 | 2.51 | 1.35 | 9.73 | 0.00 | 0.00 | 0.00 |
| 304 | piR-hsa-4450044 | 0.53 | 1.54 | 1.20 | 13.38 | 0.00 | 0.00 | 0.00 | 0.00 | 0.00 |
| 305 | piR-hsa-3741185 | 1.59 | 0.00 | 0.00 | 0.00 | 1.35 | 1.95 | 8.02 | 0.90 | 2.79 |
| 306 | piR-hsa-4144265 | 1.59 | 3.08 | 6.02 | 0.84 | 0.00 | 3.89 | 0.00 | 0.45 | 0.70 |
| 307 | piR-hsa-4198101 | 1.59 | 7.69 | 1.20 | 0.84 | 1.35 | 3.89 | 0.00 | 0.00 | 0.00 |
| 308 | piR-hsa-8117137 | 0.00 | 0.00 | 2.41 | 0.84 | 0.00 | 1.95 | 0.00 | 0.90 | 10.46 |
| 309 | piR-hsa-4100164 | 1.06 | 4.62 | 4.82 | 1.67 | 2.70 | 0.00 | 1.60 | 0.00 | 0.00 |
| 310 | piR-hsa-3280518 | 6.36 | 3.08 | 3.61 | 3.35 | 0.00 | 0.00 | 0.00 | 0.00 | 0.00 |
| 311 | piR-hsa-3710717 | 0.00 | 0.00 | 0.00 | 3.35 | 1.35 | 11.68 | 0.00 | 0.00 | 0.00 |
| 312 | piR-hsa-848451 | 0.00 | 0.00 | 0.00 | 2.51 | 4.05 | 9.73 | 0.00 | 0.00 | 0.00 |
| 313 | piR-hsa-1057272 | 0.00 | 0.00 | 0.00 | 0.00 | 1.35 | 0.00 | 11.23 | 2.26 | 1.39 |
| 314 | piR-hsa-343616 | 0.00 | 0.00 | 0.00 | 2.51 | 0.00 | 5.84 | 1.60 | 1.36 | 4.88 |
| 315 | piR-hsa-2832439 | 1.59 | 1.54 | 1.20 | 0.84 | 0.00 | 1.95 | 4.81 | 1.36 | 2.79 |
| 316 | piR-hsa-2889978 | 1.06 | 1.54 | 1.20 | 5.02 | 1.35 | 5.84 | 0.00 | 0.00 | 0.00 |
| 317 | piR-hsa-2529368 | 3.18 | 1.54 | 1.20 | 0.00 | 0.00 | 0.00 | 1.60 | 1.36 | 6.97 |
| 318 | piR-hsa-3161050 | 0.00 | 0.00 | 0.00 | 5.86 | 4.05 | 3.89 | 1.60 | 0.45 | 0.00 |
| 319 | piR-hsa-1688824 | 2.12 | 1.54 | 1.20 | 5.86 | 2.70 | 1.95 | 0.00 | 0.45 | 0.00 |
| 320 | piR-hsa-3638679 | 4.24 | 4.62 | 0.00 | 1.67 | 1.35 | 3.89 | 0.00 | 0.00 | 0.00 |
| 321 | piR-hsa-2281305 | 0.00 | 0.00 | 1.20 | 0.00 | 0.00 | 0.00 | 9.62 | 1.36 | 3.49 |
| 322 | piR-hsa-2427082 | 3.18 | 4.62 | 4.82 | 1.67 | 1.35 | 0.00 | 0.00 | 0.00 | 0.00 |
| 323 | piR-hsa-5062317 | 0.00 | 4.62 | 0.00 | 0.84 | 0.00 | 0.00 | 0.00 | 1.81 | 8.37 |
| 324 | piR-hsa-1555218 | 1.59 | 4.62 | 6.02 | 3.35 | 0.00 | 0.00 | 0.00 | 0.00 | 0.00 |
| 325 | piR-hsa-4391981_9 | 0.00 | 0.00 | 0.00 | 3.35 | 0.00 | 3.89 | 1.60 | 3.16 | 3.49 |
| 326 | piR-hsa-4403628 | 4.77 | 3.08 | 0.00 | 0.84 | 0.00 | 0.00 | 4.81 | 0.45 | 1.39 |
| 327 | piR-hsa-153942 | 1.06 | 1.54 | 1.20 | 0.00 | 0.00 | 0.00 | 8.02 | 1.36 | 2.09 |
| 328 | piR-hsa-2856544 | 4.77 | 1.54 | 2.41 | 1.67 | 0.00 | 0.00 | 3.21 | 0.90 | 0.70 |
| 329 | piR-hsa-1894768 | 1.59 | 1.54 | 1.20 | 0.00 | 0.00 | 1.95 | 3.21 | 3.61 | 2.09 |
| 330 | piR-hsa-2495779 | 2.65 | 1.54 | 1.20 | 0.00 | 4.05 | 0.00 | 1.60 | 2.71 | 1.39 |
| 331 | piR-hsa-3706918 | 0.00 | 0.00 | 0.00 | 3.35 | 0.00 | 0.00 | 6.42 | 1.81 | 3.49 |
| 332 | piR-hsa-4157592 | 2.12 | 1.54 | 1.20 | 4.18 | 4.05 | 1.95 | 0.00 | 0.00 | 0.00 |
| 333 | piR-hsa-4175186 | 0.53 | 1.54 | 9.63 | 0.00 | 1.35 | 1.95 | 0.00 | 0.00 | 0.00 |
| 334 | piR-hsa-3714350 | 0.53 | 0.00 | 0.00 | 9.20 | 1.35 | 0.00 | 3.21 | 0.00 | 0.70 |
| 335 | piR-hsa-354004 | 1.06 | 10.77 | 1.20 | 0.00 | 0.00 | 1.95 | 0.00 | 0.00 | 0.00 |
| 336 | piR-hsa-3694141 | 0.00 | 1.54 | 1.20 | 10.87 | 1.35 | 0.00 | 0.00 | 0.00 | 0.00 |
| 337 | piR-hsa-4403262 | 0.00 | 0.00 | 0.00 | 0.00 | 0.00 | 0.00 | 12.83 | 0.00 | 2.09 |
| 338 | piR-hsa-3383102 | 1.59 | 3.08 | 0.00 | 4.18 | 4.05 | 1.95 | 0.00 | 0.00 | 0.00 |
| 339 | piR-hsa-3357365 | 3.71 | 7.69 | 1.20 | 0.84 | 1.35 | 0.00 | 0.00 | 0.00 | 0.00 |
| 340 | piR-hsa-213829 | 0.53 | 4.62 | 1.20 | 8.37 | 0.00 | 0.00 | 0.00 | 0.00 | 0.00 |
| 341 | piR-hsa-7893387 | 0.53 | 0.00 | 0.00 | 2.51 | 8.09 | 1.95 | 1.60 | 0.00 | 0.00 |
| 342 | piR-hsa-235205 | 0.00 | 0.00 | 0.00 | 0.00 | 0.00 | 0.00 | 12.83 | 0.45 | 1.39 |
| 343 | piR-hsa-1416366 | 0.00 | 0.00 | 0.00 | 11.71 | 1.35 | 0.00 | 1.60 | 0.00 | 0.00 |
| 344 | piR-hsa-1998991 | 2.12 | 9.23 | 0.00 | 0.00 | 1.35 | 1.95 | 0.00 | 0.00 | 0.00 |
| 345 | piR-hsa-5124632 | 1.59 | 7.69 | 1.20 | 0.84 | 1.35 | 1.95 | 0.00 | 0.00 | 0.00 |
| 346 | piR-hsa-3365644 | 0.00 | 0.00 | 2.41 | 7.53 | 2.70 | 1.95 | 0.00 | 0.00 | 0.00 |
| 347 | piR-hsa-1886018 | 0.53 | 0.00 | 0.00 | 0.00 | 0.00 | 0.00 | 11.23 | 1.36 | 1.39 |
| 348 | piR-hsa-3657078 | 6.89 | 0.00 | 2.41 | 0.84 | 1.35 | 0.00 | 0.00 | 0.90 | 2.09 |
| 349 | piR-hsa-3104125 | 4.24 | 1.54 | 4.82 | 2.51 | 1.35 | 0.00 | 0.00 | 0.00 | 0.00 |
| 350 | piR-hsa-4378137_4 | 0.53 | 1.54 | 0.00 | 4.18 | 2.70 | 3.89 | 0.00 | 0.90 | 0.70 |
| 351 | piR-hsa-1592257 | 0.00 | 0.00 | 0.00 | 0.00 | 0.00 | 0.00 | 11.23 | 1.81 | 1.39 |
| 352 | piR-hsa-3818788 | 4.24 | 0.00 | 6.02 | 0.00 | 4.05 | 0.00 | 0.00 | 0.00 | 0.00 |
| 353 | piR-hsa-2839864 | 1.06 | 0.00 | 0.00 | 0.84 | 0.00 | 0.00 | 6.42 | 3.16 | 2.79 |
| 354 | piR-hsa-1678085 | 0.00 | 0.00 | 1.20 | 1.67 | 9.44 | 1.95 | 0.00 | 0.00 | 0.00 |

| | piRNA | psc1 | psc2 | psc3 | mpc1 | mpc2 | mpc3 | cpc1 | cpc2 | cpc3 |
|---|---|---|---|---|---|---|---|---|---|---|
| 355 | piR-hsa-4193743 | 0.00 | 0.00 | 0.00 | 0.00 | 0.00 | 0.00 | 11.23 | 0.90 | 2.09 |
| 356 | piR-hsa-2537452 | 0.00 | 0.00 | 0.00 | 14.22 | 0.00 | 0.00 | 0.00 | 0.00 | 0.00 |
| 357 | piR-hsa-4507261 | 0.00 | 1.54 | 0.00 | 0.00 | 0.00 | 0.00 | 0.00 | 2.71 | 9.76 |
| 358 | piR-hsa-1263612 | 6.89 | 0.00 | 4.82 | 0.00 | 0.00 | 0.00 | 0.00 | 0.90 | 1.39 |
| 359 | piR-hsa-778924 | 0.53 | 1.54 | 0.00 | 7.53 | 2.70 | 0.00 | 1.60 | 0.00 | 0.00 |
| 360 | piR-hsa-2165649 | 1.06 | 4.62 | 4.82 | 0.00 | 1.35 | 1.95 | 0.00 | 0.00 | 0.00 |
| 361 | piR-hsa-1274138 | 1.06 | 0.00 | 0.00 | 0.00 | 1.35 | 1.95 | 6.42 | 0.90 | 2.09 |
| 362 | piR-hsa-4111185 | 1.06 | 3.08 | 1.20 | 7.53 | 0.00 | 0.00 | 0.00 | 0.00 | 0.70 |
| 363 | piR-hsa-3693411 | 0.00 | 0.00 | 0.00 | 5.86 | 2.70 | 1.95 | 1.60 | 0.00 | 1.39 |
| 364 | piR-hsa-1917013 | 0.53 | 3.08 | 0.00 | 0.00 | 0.00 | 0.00 | 6.42 | 2.71 | 0.70 |
| 365 | piR-hsa-4467055_8 | 0.00 | 0.00 | 2.41 | 5.86 | 0.00 | 3.89 | 0.00 | 0.45 | 0.70 |
| 366 | piR-hsa-4402141 | 0.00 | 0.00 | 0.00 | 0.00 | 0.00 | 0.00 | 11.23 | 1.36 | 0.70 |
| 367 | piR-hsa-942735 | 0.00 | 0.00 | 0.00 | 0.00 | 0.00 | 0.00 | 11.23 | 1.36 | 0.70 |
| 368 | piR-hsa-1574545 | 0.00 | 0.00 | 0.00 | 0.84 | 0.00 | 0.00 | 9.62 | 0.00 | 2.79 |
| 369 | piR-hsa-2161668 | 0.00 | 0.00 | 1.20 | 4.18 | 5.39 | 1.95 | 0.00 | 0.45 | 0.00 |
| 370 | piR-hsa-771001 | 0.53 | 0.00 | 0.00 | 9.20 | 1.35 | 0.00 | 1.60 | 0.45 | 0.00 |
| 371 | piR-hsa-2502273 | 3.71 | 0.00 | 4.82 | 2.51 | 0.00 | 1.95 | 0.00 | 0.00 | 0.00 |
| 372 | piR-hsa-77303 | 2.65 | 1.54 | 6.02 | 0.00 | 1.35 | 0.00 | 0.00 | 0.00 | 1.39 |
| 373 | piR-hsa-2853562 | 0.00 | 0.00 | 1.20 | 8.37 | 1.35 | 1.95 | 0.00 | 0.00 | 0.00 |
| 374 | piR-hsa-2851130 | 1.59 | 0.00 | 0.00 | 0.00 | 0.00 | 0.00 | 4.81 | 2.26 | 4.18 |
| 375 | piR-hsa-2538984 | 7.42 | 0.00 | 0.00 | 0.84 | 1.35 | 0.00 | 1.60 | 0.90 | 0.70 |
| 376 | piR-hsa-748358_5 | 0.00 | 0.00 | 0.00 | 0.00 | 0.00 | 0.00 | 9.62 | 3.16 | 0.00 |
| 377 | piR-hsa-2400882 | 0.53 | 0.00 | 0.00 | 2.51 | 5.39 | 3.89 | 0.00 | 0.45 | 0.00 |
| 378 | piR-hsa-3610092 | 5.30 | 1.54 | 2.41 | 0.00 | 1.35 | 1.95 | 0.00 | 0.00 | 0.00 |
| 379 | piR-hsa-727158 | 5.83 | 1.54 | 3.61 | 0.84 | 0.00 | 0.00 | 0.00 | 0.00 | 0.70 |
| 380 | piR-hsa-1530800 | 0.00 | 0.00 | 0.00 | 5.86 | 2.70 | 3.89 | 0.00 | 0.00 | 0.00 |
| 381 | piR-hsa-1603046 | 0.00 | 0.00 | 0.00 | 2.51 | 4.05 | 5.84 | 0.00 | 0.00 | 0.00 |
| 382 | piR-hsa-2518229 | 0.00 | 1.54 | 0.00 | 0.00 | 1.35 | 0.00 | 3.21 | 1.81 | 4.18 |
| 383 | piR-hsa-3642754 | 4.77 | 1.54 | 4.82 | 0.84 | 0.00 | 0.00 | 0.00 | 0.00 | 0.00 |
| 384 | piR-hsa-5039329_2 | 0.00 | 0.00 | 0.00 | 0.00 | 0.00 | 0.00 | 9.62 | 0.90 | 1.39 |
| 385 | piR-hsa-4120912 | 0.53 | 0.00 | 0.00 | 3.35 | 5.39 | 1.95 | 0.00 | 0.00 | 0.70 |
| 386 | piR-hsa-3263398 | 5.83 | 3.08 | 0.00 | 0.00 | 0.00 | 0.00 | 0.00 | 2.26 | 0.70 |
| 387 | piR-hsa-2484024 | 0.00 | 0.00 | 0.00 | 0.84 | 0.00 | 0.00 | 8.02 | 0.90 | 2.09 |
| 388 | piR-hsa-4388882 | 0.53 | 1.54 | 0.00 | 8.37 | 1.35 | 0.00 | 0.00 | 0.00 | 0.00 |
| 389 | piR-hsa-2499630 | 0.53 | 0.00 | 0.00 | 0.00 | 0.00 | 0.00 | 8.02 | 0.90 | 2.09 |
| 390 | piR-hsa-2515298 | 4.24 | 1.54 | 0.00 | 0.00 | 0.00 | 0.00 | 3.21 | 0.45 | 2.09 |
| 391 | piR-hsa-65029 | 2.12 | 1.54 | 4.82 | 1.67 | 1.35 | 0.00 | 0.00 | 0.00 | 0.00 |
| 392 | piR-hsa-271367 | 3.71 | 3.08 | 2.41 | 0.84 | 1.35 | 0.00 | 0.00 | 0.00 | 0.00 |
| 393 | piR-hsa-2501382 | 1.06 | 0.00 | 1.20 | 0.84 | 0.00 | 0.00 | 3.21 | 2.26 | 2.79 |
| 394 | piR-hsa-3345389 | 0.00 | 0.00 | 0.00 | 3.35 | 4.05 | 3.89 | 0.00 | 0.00 | 0.00 |
| 395 | piR-hsa-1881756 | 0.00 | 0.00 | 0.00 | 0.00 | 1.35 | 0.00 | 6.42 | 0.00 | 3.49 |
| 396 | piR-hsa-1593686 | 3.71 | 3.08 | 1.20 | 0.84 | 0.00 | 1.95 | 0.00 | 0.45 | 0.00 |
| 397 | piR-hsa-4359698 | 2.12 | 4.62 | 2.41 | 0.00 | 0.00 | 1.95 | 0.00 | 0.00 | 0.00 |
| 398 | piR-hsa-1866304 | 3.18 | 3.08 | 4.82 | 0.00 | 0.00 | 0.00 | 0.00 | 0.00 | 0.00 |
| 399 | piR-hsa-1550295 | 2.12 | 7.69 | 1.20 | 0.00 | 0.00 | 0.00 | 0.00 | 0.00 | 0.00 |
| 400 | piR-hsa-550050 | 0.00 | 0.00 | 0.00 | 0.00 | 0.00 | 0.00 | 6.42 | 3.16 | 1.39 |
| 401 | piR-hsa-76694 | 3.71 | 0.00 | 7.23 | 0.00 | 0.00 | 0.00 | 0.00 | 0.00 | 0.00 |
| 402 | piR-hsa-2491463 | 4.24 | 3.08 | 0.00 | 0.00 | 0.00 | 1.95 | 0.00 | 0.90 | 0.70 |
| 403 | piR-hsa-2534860 | 1.06 | 0.00 | 0.00 | 0.00 | 1.35 | 3.89 | 0.00 | 3.16 | 1.39 |
| 404 | piR-hsa-2715002 | 0.00 | 0.00 | 0.00 | 0.00 | 0.00 | 0.00 | 8.02 | 1.36 | 1.39 |
| 405 | piR-hsa-2511205 | 0.53 | 0.00 | 0.00 | 0.84 | 0.00 | 0.00 | 3.21 | 4.07 | 2.09 |
| 406 | piR-hsa-4271500 | 0.53 | 1.54 | 0.00 | 6.69 | 0.00 | 1.95 | 0.00 | 0.00 | 0.00 |
| 407 | piR-hsa-4150185 | 1.06 | 0.00 | 1.20 | 8.37 | 0.00 | 0.00 | 0.00 | 0.00 | 0.00 |
| 408 | piR-hsa-1943369 | 0.00 | 0.00 | 0.00 | 5.02 | 5.39 | 0.00 | 0.00 | 0.00 | 0.00 |
| 409 | piR-hsa-2447480 | 1.59 | 3.08 | 4.82 | 0.84 | 0.00 | 0.00 | 0.00 | 0.00 | 0.00 |
| 410 | piR-hsa-1416960 | 2.65 | 4.62 | 1.20 | 1.67 | 0.00 | 0.00 | 0.00 | 0.00 | 0.00 |
| 411 | piR-hsa-699024 | 0.00 | 0.00 | 1.20 | 0.00 | 0.00 | 0.00 | 6.42 | 1.81 | 0.70 |
| 412 | piR-hsa-6764147 | 0.00 | 0.00 | 1.20 | 0.84 | 0.00 | 0.00 | 0.00 | 1.81 | 6.28 |
| 413 | piR-hsa-644441_3 | 0.53 | 0.00 | 0.00 | 0.00 | 0.00 | 0.00 | 6.42 | 3.16 | 0.00 |

| | piRNA | psc1 | psc2 | psc3 | mpc1 | mpc2 | mpc3 | cpc1 | cpc2 | cpc3 |
|---|---|---|---|---|---|---|---|---|---|---|
| 414 | piR-hsa-2500917 | 0.53 | 0.00 | 1.20 | 0.00 | 0.00 | 0.00 | 4.81 | 1.36 | 2.09 |
| 415 | piR-hsa-1160813 | 0.00 | 0.00 | 0.00 | 6.69 | 1.35 | 1.95 | 0.00 | 0.00 | 0.00 |
| 416 | piR-hsa-628215 | 0.00 | 0.00 | 0.00 | 0.00 | 0.00 | 0.00 | 6.42 | 0.00 | 3.49 |
| 417 | piR-hsa-5313694 | 0.00 | 0.00 | 0.00 | 0.00 | 0.00 | 0.00 | 4.81 | 4.97 | 0.00 |
| 418 | piR-hsa-778446 | 0.53 | 0.00 | 0.00 | 0.00 | 0.00 | 0.00 | 1.60 | 1.36 | 6.28 |
| 419 | piR-hsa-313568 | 0.00 | 0.00 | 0.00 | 8.37 | 1.35 | 0.00 | 0.00 | 0.00 | 0.00 |
| 420 | piR-hsa-3153428 | 2.65 | 4.62 | 2.41 | 0.00 | 0.00 | 0.00 | 0.00 | 0.00 | 0.00 |
| 421 | piR-hsa-2365951 | 0.00 | 0.00 | 0.00 | 5.86 | 1.35 | 1.95 | 0.00 | 0.45 | 0.00 |
| 422 | piR-hsa-4398177 | 0.00 | 0.00 | 0.00 | 8.37 | 0.00 | 0.00 | 0.00 | 0.45 | 0.70 |
| 423 | piR-hsa-5884872 | 1.59 | 3.08 | 4.82 | 0.00 | 0.00 | 0.00 | 0.00 | 0.00 | 0.00 |
| 424 | piR-hsa-851842 | 3.71 | 0.00 | 3.61 | 0.00 | 0.00 | 1.95 | 0.00 | 0.00 | 0.00 |
| 425 | piR-hsa-4175241 | 0.53 | 0.00 | 1.20 | 7.53 | 0.00 | 0.00 | 0.00 | 0.00 | 0.00 |
| 426 | piR-hsa-612387 | 3.71 | 3.08 | 2.41 | 0.00 | 0.00 | 0.00 | 0.00 | 0.00 | 0.00 |
| 427 | piR-hsa-1551388 | 0.00 | 0.00 | 0.00 | 0.00 | 0.00 | 0.00 | 4.81 | 2.26 | 2.09 |
| 428 | piR-hsa-4049552 | 0.00 | 0.00 | 0.00 | 0.00 | 1.35 | 0.00 | 3.21 | 1.81 | 2.79 |
| 429 | piR-hsa-42060 | 1.06 | 0.00 | 0.00 | 6.69 | 1.35 | 0.00 | 0.00 | 0.00 | 0.00 |
| 430 | piR-hsa-338024 | 3.18 | 1.54 | 2.41 | 0.00 | 0.00 | 0.00 | 0.00 | 0.45 | 1.39 |
| 431 | piR-hsa-2030201 | 3.71 | 1.54 | 3.61 | 0.00 | 0.00 | 0.00 | 0.00 | 0.00 | 0.00 |
| 432 | piR-hsa-4135859 | 0.00 | 0.00 | 0.00 | 0.84 | 0.00 | 0.00 | 3.21 | 4.07 | 0.70 |
| 433 | piR-hsa-1692624 | 0.00 | 0.00 | 0.00 | 6.69 | 0.00 | 1.95 | 0.00 | 0.00 | 0.00 |
| 434 | piR-hsa-1534285 | 2.12 | 1.54 | 4.82 | 0.00 | 0.00 | 0.00 | 0.00 | 0.00 | 0.00 |
| 435 | piR-hsa-2832794 | 0.00 | 0.00 | 0.00 | 0.00 | 0.00 | 0.00 | 4.81 | 2.26 | 1.39 |
| 436 | piR-hsa-1370047 | 0.00 | 0.00 | 0.00 | 0.84 | 0.00 | 0.00 | 0.00 | 5.87 | 1.39 |
| 437 | piR-hsa-632933 | 0.00 | 0.00 | 0.00 | 0.00 | 1.35 | 0.00 | 1.60 | 2.26 | 2.79 |
| 438 | piR-hsa-2069180 | 4.24 | 0.00 | 1.20 | 0.84 | 1.35 | 0.00 | 0.00 | 0.00 | 0.00 |
| 439 | piR-hsa-1524373 | 0.00 | 0.00 | 0.00 | 0.00 | 0.00 | 0.00 | 1.60 | 0.90 | 4.88 |
| 440 | piR-hsa-2487038_7 | 0.00 | 0.00 | 0.00 | 0.00 | 0.00 | 0.00 | 0.00 | 1.81 | 5.58 |
| 441 | piR-hsa-5568482 | 0.00 | 0.00 | 0.00 | 0.00 | 0.00 | 0.00 | 0.00 | 3.61 | 3.49 |
| 442 | piR-hsa-4346643 | 0.00 | 0.00 | 0.00 | 0.84 | 0.00 | 0.00 | 0.00 | 4.07 | 2.09 |
| 443 | piR-hsa-7313613 | 0.53 | 1.54 | 0.00 | 0.00 | 0.00 | 0.00 | 0.00 | 3.61 | 0.70 |
| 444 | piR-hsa-4398796 | 4.77 | 0.00 | 0.00 | 0.00 | 0.00 | 0.00 | 0.00 | 1.36 | 0.00 |
| 445 | piR-hsa-2369565 | 0.00 | 0.00 | 0.00 | 0.00 | 0.00 | 0.00 | 0.00 | 0.90 | 4.88 |
| 446 | piR-hsa-7098338 | 0.00 | 0.00 | 0.00 | 0.00 | 0.00 | 0.00 | 1.60 | 3.16 | 0.70 |
| 447 | piR-hsa-7670531_2 | 0.00 | 0.00 | 0.00 | 0.00 | 0.00 | 0.00 | 0.00 | 3.61 | 1.39 |

Table S2: Normalized counts for the 447 identified piRNAs arranged in descending order expression (row mean). Differentially regulated piRNAs in CPC are indicated in light blue shading (downregulated) and light red shading (upregulated).

| | cluster1 | cluster2 | cluster3 | cluster4 | cluster5 | cluster6 | cluster7 | cluster8 |
|---|---|---|---|---|---|---|---|---|
| 1 | piR-hsa-108574 | piR-hsa-1001021 | piR-hsa-118348 | piR-hsa-114666 | piR-hsa-1233052 | piR-hsa-1904126 | piR-hsa-1205256 | piR-hsa-151466 |
| 2 | piR-hsa-132896 | piR-hsa-1277994 | piR-hsa-1263612 | piR-hsa-1389062 | piR-hsa-1870459 | piR-hsa-1916259 | piR-hsa-1429070 | piR-hsa-1548068 |
| 3 | piR-hsa-137136 | piR-hsa-147461 | piR-hsa-1291516 | piR-hsa-1463989 | piR-hsa-1870588 | piR-hsa-359160_2 | piR-hsa-1690788 | piR-hsa-1875212 |
| 4 | piR-hsa-1399886 | piR-hsa-151249 | piR-hsa-1332287 | piR-hsa-151136 | piR-hsa-1872235 | piR-hsa-3732777 | piR-hsa-1773241 | piR-hsa-1876265 |
| 5 | piR-hsa-1492262 | piR-hsa-1872085_4 | piR-hsa-1376916 | piR-hsa-163695 | piR-hsa-1890632 | piR-hsa-1862211 | piR-hsa-1903779 | piR-hsa-1883972 |
| 6 | piR-hsa-1843231 | piR-hsa-2482189 | piR-hsa-1409954 | piR-hsa-1647694 | piR-hsa-1900529 | piR-hsa-1872463 | piR-hsa-2151268 | piR-hsa-1908839 |
| 7 | piR-hsa-2090890 | piR-hsa-2646470 | piR-hsa-1416960 | piR-hsa-169217 | piR-hsa-1919455 | piR-hsa-6744266 | piR-hsa-2252211 | piR-hsa-1919272 |
| 8 | piR-hsa-2172087 | piR-hsa-611204 | piR-hsa-1528884 | piR-hsa-1901970 | piR-hsa-1927965 | | piR-hsa-2395910 | piR-hsa-1920687 |
| 9 | piR-hsa-2333057 | piR-hsa-76848 | piR-hsa-1534285 | piR-hsa-1988800 | piR-hsa-1941780 | | piR-hsa-2398570 | piR-hsa-1923208 |
| 10 | piR-hsa-2536290 | piR-hsa-1160813 | piR-hsa-1550295 | piR-hsa-20628 | piR-hsa-2477264 | | piR-hsa-2513278 | piR-hsa-2490897 |
| 11 | piR-hsa-2856604 | piR-hsa-1229611 | piR-hsa-1555218 | piR-hsa-214132 | piR-hsa-2479371 | | piR-hsa-255984 | piR-hsa-2521457 |
| 12 | piR-hsa-2882083 | piR-hsa-1256360 | piR-hsa-1593307 | piR-hsa-2152778 | piR-hsa-2481097 | | piR-hsa-2565910 | piR-hsa-2526525 |
| 13 | piR-hsa-298158 | piR-hsa-1259933 | piR-hsa-1593686 | piR-hsa-2208850 | piR-hsa-2494226 | | piR-hsa-2615134 | piR-hsa-2539762 |
| 14 | piR-hsa-3021684 | piR-hsa-1302552 | piR-hsa-1595580 | piR-hsa-2286229 | piR-hsa-2515454 | | piR-hsa-2863156 | piR-hsa-316012 |
| 15 | piR-hsa-3119265 | piR-hsa-1303811 | piR-hsa-1607096 | piR-hsa-2299252 | piR-hsa-2530015 | | piR-hsa-3232943 | piR-hsa-343382 |
| 16 | piR-hsa-3546008 | piR-hsa-1340768 | piR-hsa-1632961 | piR-hsa-2346976 | piR-hsa-2831593 | | piR-hsa-3352181 | piR-hsa-3634065 |
| 17 | piR-hsa-5077723 | piR-hsa-1416366 | piR-hsa-163499 | piR-hsa-2780538 | piR-hsa-2832647 | | piR-hsa-3683883 | piR-hsa-3658275 |
| 18 | piR-hsa-6913457 | piR-hsa-1425899 | piR-hsa-1686806 | piR-hsa-2844156 | piR-hsa-2840936 | | piR-hsa-389007 | piR-hsa-3658742 |
| 19 | piR-hsa-721859_9 | piR-hsa-1434629 | piR-hsa-1726249 | piR-hsa-3265318 | piR-hsa-346271 | | piR-hsa-4028152 | piR-hsa-3739406 |
| 20 | | piR-hsa-1481120 | piR-hsa-1748898 | piR-hsa-3660133 | piR-hsa-3623001 | | piR-hsa-4381848_2 | piR-hsa-4460706 |
| 21 | | piR-hsa-1530800 | piR-hsa-1798104 | piR-hsa-3817390 | piR-hsa-368987 | | piR-hsa-4472891 | piR-hsa-1576285 |
| 22 | | piR-hsa-1531418 | piR-hsa-1866304 | piR-hsa-3875265 | piR-hsa-3735787 | | piR-hsa-724912 | piR-hsa-785969 |
| 23 | | piR-hsa-1557538 | piR-hsa-1875155 | piR-hsa-3916570 | piR-hsa-768321 | | piR-hsa-7765828 | piR-hsa-1057272 |
| 24 | | piR-hsa-1585146 | piR-hsa-1929067 | piR-hsa-4078407 | piR-hsa-771714_3 | | | piR-hsa-1242358 |
| 25 | | piR-hsa-1603046 | piR-hsa-1998991 | piR-hsa-4303719 | piR-hsa-7892960 | | | piR-hsa-1259653 |
| 26 | | piR-hsa-160969 | piR-hsa-2027369 | piR-hsa-5982715 | | | | piR-hsa-1274138 |
| 27 | | piR-hsa-161264_3 | piR-hsa-2030201 | piR-hsa-6245615 | | | | piR-hsa-1284504 |
| 28 | | piR-hsa-1678085 | piR-hsa-2069180 | piR-hsa-783698 | | | | piR-hsa-1296118 |
| 29 | | piR-hsa-1688824 | piR-hsa-2090264 | | | | | piR-hsa-1370047 |
| 30 | | piR-hsa-1692624 | piR-hsa-2137611 | | | | | piR-hsa-144277_2 |
| 31 | | piR-hsa-1706026 | piR-hsa-2148238 | | | | | piR-hsa-147696 |
| 32 | | piR-hsa-1708978 | piR-hsa-2165649 | | | | | piR-hsa-1524373 |
| 33 | | piR-hsa-1756618 | piR-hsa-2213434 | | | | | piR-hsa-153942 |
| 34 | | piR-hsa-1883893 | piR-hsa-2230204 | | | | | piR-hsa-1551388 |
| 35 | | piR-hsa-1905329 | piR-hsa-2240007 | | | | | piR-hsa-1574545 |
| 36 | | piR-hsa-1909905 | piR-hsa-2248086 | | | | | piR-hsa-1592257 |
| 37 | | piR-hsa-1921551 | piR-hsa-2268195 | | | | | piR-hsa-1696540 |
| 38 | | piR-hsa-1922210 | piR-hsa-2308163 | | | | | piR-hsa-1707103 |

| | cluster1 | cluster2 | cluster3 | cluster4 | cluster5 | cluster6 | cluster7 | cluster8 |
|---|---|---|---|---|---|---|---|---|
| 39 | | piR-hsa-1933276 | piR-hsa-2319750 | | | | | piR-hsa-1881756 |
| 40 | | piR-hsa-1939085 | piR-hsa-2351941 | | | | | piR-hsa-1882039 |
| 41 | | piR-hsa-1941637 | piR-hsa-2353109 | | | | | piR-hsa-1886018 |
| 42 | | piR-hsa-1943369 | piR-hsa-2398119 | | | | | piR-hsa-1894768 |
| 43 | | piR-hsa-211123 | piR-hsa-2413094 | | | | | piR-hsa-1905680 |
| 44 | | piR-hsa-213829 | piR-hsa-2426792 | | | | | piR-hsa-1912443 |
| 45 | | piR-hsa-2161668 | piR-hsa-2427082 | | | | | piR-hsa-1917013 |
| 46 | | piR-hsa-21839 | piR-hsa-2436454 | | | | | piR-hsa-1917139 |
| 47 | | piR-hsa-2209630 | piR-hsa-2447480 | | | | | piR-hsa-1921188 |
| 48 | | piR-hsa-2220917 | piR-hsa-2491463 | | | | | piR-hsa-1927627 |
| 49 | | piR-hsa-2253283 | piR-hsa-2497478 | | | | | piR-hsa-1938524 |
| 50 | | piR-hsa-229786 | piR-hsa-2502273 | | | | | piR-hsa-2042088 |
| 51 | | piR-hsa-2365951 | piR-hsa-2503702 | | | | | piR-hsa-2072163 |
| 52 | | piR-hsa-237221 | piR-hsa-2515298 | | | | | piR-hsa-2281305 |
| 53 | | piR-hsa-2400882 | piR-hsa-2519215 | | | | | piR-hsa-235205 |
| 54 | | piR-hsa-2423519 | piR-hsa-2538984 | | | | | piR-hsa-2369565 |
| 55 | | piR-hsa-2450089_2 | piR-hsa-271367 | | | | | piR-hsa-2425220 |
| 56 | | piR-hsa-2464166 | piR-hsa-2856544 | | | | | piR-hsa-2478880_2 |
| 57 | | piR-hsa-2489909 | piR-hsa-2989729 | | | | | piR-hsa-2484024 |
| 58 | | piR-hsa-2505515 | piR-hsa-3104125 | | | | | piR-hsa-2487038_7 |
| 59 | | piR-hsa-2525461 | piR-hsa-3136454 | | | | | piR-hsa-2490287 |
| 60 | | piR-hsa-2537452 | piR-hsa-3153428 | | | | | piR-hsa-2490509 |
| 61 | | piR-hsa-2589139 | piR-hsa-315964 | | | | | piR-hsa-2495779 |
| 62 | | piR-hsa-2670375 | piR-hsa-3177742 | | | | | piR-hsa-2499630 |
| 63 | | piR-hsa-2853562 | piR-hsa-3231825 | | | | | piR-hsa-2500917 |
| 64 | | piR-hsa-2889978 | piR-hsa-3263398 | | | | | piR-hsa-2501382 |
| 65 | | piR-hsa-307961 | piR-hsa-3280518 | | | | | piR-hsa-2511205 |
| 66 | | piR-hsa-313568 | piR-hsa-333507 | | | | | piR-hsa-2518229 |
| 67 | | piR-hsa-3161050 | piR-hsa-3357365 | | | | | piR-hsa-2529368 |
| 68 | | piR-hsa-3345389 | piR-hsa-338024 | | | | | piR-hsa-2534860 |
| 69 | | piR-hsa-3365644 | piR-hsa-3513154 | | | | | piR-hsa-2542835 |
| 70 | | piR-hsa-3383102 | piR-hsa-3527815 | | | | | piR-hsa-2592846 |
| 71 | | piR-hsa-362913 | piR-hsa-354004 | | | | | piR-hsa-2715002 |
| 72 | | piR-hsa-3634880 | piR-hsa-3558751 | | | | | piR-hsa-2742244 |
| 73 | | piR-hsa-3693411 | piR-hsa-3610092 | | | | | piR-hsa-2827579 |
| 74 | | piR-hsa-3694141 | piR-hsa-3638679 | | | | | piR-hsa-2829413 |
| 75 | | piR-hsa-3710717 | piR-hsa-3642754 | | | | | piR-hsa-2829712 |
| 76 | | piR-hsa-3714350 | piR-hsa-3657078 | | | | | piR-hsa-2832439 |

| | cluster1 | cluster2 | cluster3 | cluster4 | cluster5 | cluster6 | cluster7 | cluster8 |
|---|---|---|---|---|---|---|---|---|
| 77 | | piR-hsa-374600 | piR-hsa-3674332 | | | | | piR-hsa-2832794 |
| 78 | | piR-hsa-3776081 | piR-hsa-3732088 | | | | | piR-hsa-2839864 |
| 79 | | piR-hsa-3829948 | piR-hsa-3807498 | | | | | piR-hsa-2851130 |
| 80 | | piR-hsa-3839126 | piR-hsa-3818788 | | | | | piR-hsa-343616 |
| 81 | | piR-hsa-3842249 | piR-hsa-3944431 | | | | | piR-hsa-363100_2 |
| 82 | | piR-hsa-3978322 | piR-hsa-3974794 | | | | | piR-hsa-368381 |
| 83 | | piR-hsa-4111185 | piR-hsa-4020841 | | | | | piR-hsa-3706918 |
| 84 | | piR-hsa-4120912 | piR-hsa-4030155 | | | | | piR-hsa-3718263_3 |
| 85 | | piR-hsa-4131663 | piR-hsa-4053516 | | | | | piR-hsa-3741185 |
| 86 | | piR-hsa-4150185 | piR-hsa-4091280 | | | | | piR-hsa-4049552 |
| 87 | | piR-hsa-4157592 | piR-hsa-4100164 | | | | | piR-hsa-4135859 |
| 88 | | piR-hsa-4175241 | piR-hsa-4110708 | | | | | piR-hsa-4193743 |
| 89 | | piR-hsa-4202081 | piR-hsa-4144265 | | | | | piR-hsa-4346643 |
| 90 | | piR-hsa-42060 | piR-hsa-4175186 | | | | | piR-hsa-4387218 |
| 91 | | piR-hsa-4271500 | piR-hsa-4178299 | | | | | piR-hsa-4391981_9 |
| 92 | | piR-hsa-4378137_4 | piR-hsa-4198101 | | | | | piR-hsa-4402141 |
| 93 | | piR-hsa-4388882 | piR-hsa-4322932 | | | | | piR-hsa-4403262 |
| 94 | | piR-hsa-4397384 | piR-hsa-4359698 | | | | | piR-hsa-4424378 |
| 95 | | piR-hsa-4398177 | piR-hsa-4379982 | | | | | piR-hsa-4507261 |
| 96 | | piR-hsa-4408495 | piR-hsa-4398796 | | | | | piR-hsa-5039329_2 |
| 97 | | piR-hsa-4416099_9 | piR-hsa-4403577 | | | | | piR-hsa-5062317 |
| 98 | | piR-hsa-4450044 | piR-hsa-4403628 | | | | | piR-hsa-5313694 |
| 99 | | piR-hsa-4467055_8 | piR-hsa-508592 | | | | | piR-hsa-550050 |
| 100 | | piR-hsa-623353 | piR-hsa-5124632 | | | | | piR-hsa-5568482 |
| 101 | | piR-hsa-669874 | piR-hsa-5411637 | | | | | piR-hsa-5996985 |
| 102 | | piR-hsa-7106256 | piR-hsa-58291 | | | | | piR-hsa-628215 |
| 103 | | piR-hsa-7308134 | piR-hsa-5884872 | | | | | piR-hsa-632933 |
| 104 | | piR-hsa-745484 | piR-hsa-612387 | | | | | piR-hsa-642866 |
| 105 | | piR-hsa-7544198 | piR-hsa-645846 | | | | | piR-hsa-644441_3 |
| 106 | | piR-hsa-771001 | piR-hsa-6482184 | | | | | piR-hsa-665910 |
| 107 | | piR-hsa-772699 | piR-hsa-65029 | | | | | piR-hsa-6764147 |
| 108 | | piR-hsa-7760463 | piR-hsa-67957 | | | | | piR-hsa-699024 |
| 109 | | piR-hsa-778924 | piR-hsa-727158 | | | | | piR-hsa-7098338 |
| 110 | | piR-hsa-7833890 | piR-hsa-76694 | | | | | piR-hsa-728085 |
| 111 | | piR-hsa-7893387 | piR-hsa-77303 | | | | | piR-hsa-7313613 |
| 112 | | piR-hsa-848451 | piR-hsa-8270846 | | | | | piR-hsa-748358_5 |
| 113 | | piR-hsa-97458 | piR-hsa-851842 | | | | | piR-hsa-753191 |
| 114 | | | | | | | | piR-hsa-7670531_2 |

| | cluster1 | cluster2 | cluster3 | cluster4 | cluster5 | cluster6 | cluster7 | cluster8 |
|---|---|---|---|---|---|---|---|---|
| 115 | | | | | | | | piR-hsa-778446 |
| 116 | | | | | | | | piR-hsa-7821967_3 |
| 117 | | | | | | | | piR-hsa-8117137 |
| 118 | | | | | | | | piR-hsa-834074 |
| 119 | | | | | | | | piR-hsa-942735 |

Table S3: PiRNAs included in Expression Clusters.
